# Genome-wide Identification of the Laccase Gene Family in White Jute (*Corchorus capsularis*): Potential Targets for Lignin Engineering in Bast Fibre

**DOI:** 10.1101/2024.07.17.603856

**Authors:** Subhadarshini Parida, Deepak Kumar Jha, Khushbu Kumari, Seema Pradhan, Nrisingha Dey, Shuvobrata Majumder

**Author notes:** Equality contributed (Subhadarshini Parida, Deepak Kumar Jha).

## Abstract

Jute (*Corchorus* spp.) is an important industrial bast fibre crop valued for its lignocellulosic fibres, yet the molecular basis of fibre lignification remains unexplored. Laccase (EC 1.10.3.2) is a key enzyme catalysing the final steps of lignin polymerisation. A genome-wide analysis of white jute (*Corchorus capsularis*) identified 32 putative laccase genes (*CcaLAC*s) that were phylogenetically grouped into six clades. Expression profiling revealed predominant expression in phloem tissue (16 genes), followed by leaf (8) and xylem/root tissues (4). Several *CcaLAC*s showed progressive upregulation from early growth to harvest stages. Homology with *Arabidopsis* laccases highlighted candidate genes involved in lignification, which were further supported by transcriptomic and qRT-PCR analyses. Notably, key *CcaLAC*s showed significantly reduced expression in *dlpf* (deficient lignified phloem fibre), a low-lignin white jute mutant. *CcaLAC* expression was also responsive to abiotic stresses, including abscisic acid and copper. MicroRNA target prediction identified Ath-miR397a and Ath-miR397b as potential regulators of multiple *CcaLAC*s. Structural and subcellular analyses revealed conserved motifs, transmembrane domains, and diverse cellular localisation. Gene ontology analysis linked *CcaLAC*s to lignin and phenylpropanoid biosynthesis. Among them, *CcaLAC28* and *CcaLAC32* emerged as strong candidate genes associated with phloem fibre lignification, representing promising targets for future functional validation towards developing low-lignin jute varieties.

## **1.** Introduction

Laccases (LACs) are multicopper oxidases widely present in plants, fungi, and bacteria. In plants, they catalyse the oxidation of phenolic substrates during lignin polymerisation, reducing molecular oxygen to water (**Berthet et al., 2012**). Their involvement in secondary cell wall formation and lignification has been demonstrated in tree species such as loblolly pine (*Pinus taeda*) and sycamore maple (*Acer pseudoplatanus*), as well as in model plant systems. However, the *LAC* gene family remains largely unexplored in jute (*Corchorus* spp.), a major lignocellulosic fibre crop.

Jute bast fibres are strong, biodegradable and widely used in packaging, textiles, paper and composite applications. Their comparatively high lignin content (12–26%) contributes to fibre coarseness and limits value-added applications relative to low-lignin flax (∼2%) and lignin-free cotton (**Majumder et al., 2020a**). Improving fibre quality, therefore, requires a better understanding of lignification and its regulatory genes.

The reference genomes of white jute (*Corchorus capsularis*) and tossa jute (*C. olitorius*) have enabled genome-wide identification of lignin-related gene families (**Zhang et al., 2021**). For example, six *CCoAOMT* genes were identified as contributors to lignification in jute. Laccases (*LAC*) and peroxidases (*PRX*) jointly catalyse monolignol polymerisation during secondary cell wall formation, yet *LAC* genes have received less attention in jute compared to *PRX* genes.

Genome-wide surveys report medium-to-large *LAC* gene families across plant lineages, including 17 putative *LAC* genes in *Arabidopsis thaliana*, 30 in rice (*Oryza sativa*), 49 in black cottonwood (*Populus trichocarpa*), 54 in flooded gum (*Eucalyptus grandis*), 45 in flax (*Linum usitatissimum*), 84 in cotton (*Gossypium hirsutum*), 22 in maize (*Zea mays*), 156 in wheat (*Triticum aestivum*) and 46 in tossa jute (*Corchorus olitorius*) (**Jha et al., 2025**). Functional studies showed that *AtLAC4* and *AtLAC17* are essential for lignification in *Arabidopsis* stems (**Berthet et al., 2011**), and overexpression of *GaLAC1* increased lignification in poplar (**Wang et al., 2008**).

Mutant cultivars play a crucial role in the functional validation of genes. In model plants such as *A. thaliana*, *LAC* gene-specific knockout mutants are available for study. However, such resources are not available for crops like jute. Fortunately, an X-ray-irradiated mutant white jute line, *dlpf* (deficient lignified phloem fibre; accession CMU 013, INGR No. 04107), was available (**Sengupta and Palit, 2004**). Compared with its parental cultivar JRC212, *dlpf* phloem fibre contains approximately 8.5% lignin versus 18.3% in the wild type (∼50% reduction), alongside a compensatory increase in cellulose content (∼66.5% versus ∼50.9%). The mutant is phenotypically distinguished by slower growth, shorter internodes, and undulated stem, petiole, and leaf-vein morphology, while showing leaf photosynthetic rates comparable to those of the wild type, indicating that the lignin deficiency is not accompanied by a general photosynthetic defect (**Sengupta and Palit, 2004**). Transcriptomic analysis of the *dlpf* mutant and JRC212 control jute revealed lower expression of multiple genes in the phenylpropanoid pathway, including the lignin pathway, suggesting a regulatory role for laccases in bast fibre lignification (**Chakraborty et al., 2015**).

Laccases also participate in stress responses in plants, such as desiccation, freezing, and high salinity, often through ABA-mediated signalling linked to lignin deposition (**Vishwakarma et al., 2017**). However, stress-induced *LAC* gene expression has not been examined in jute. The *in silico* study of upstream cis-acting elements of genes provides insights into the stress conditions or signalling pathways that regulate their expression. The presence of stress-, hormone- and development-related cis-elements may therefore provide functional clues.

The objective of this study is to identify and characterise the *LAC* gene family in white jute (*C. capsularis*), and to analyse its expression across tissues and developmental stages, and under copper stress and ABA treatment, to infer its potential roles in fibre lignification and stress adaptation.

The genome-wide identification of *LAC* gene families has been done previously in other crops; this study is distinguished by its integration of genome-wide discovery with mutant-based expression validation and stress-responsiveness profiling. This approach has not previously been applied to any jute LAC family, enabling prioritisation of specific candidate genes rather than a purely descriptive inventory. These analyses will help prioritise candidate *CcaLAC* genes for future functional validation and molecular breeding to improve jute fibre quality.

## 2. Materials and Methods

### 2.1. Identification of Laccase Genes in White Jute and Sequence Analysis

The white jute genomic DNA, coding sequences (CDS), and protein sequences were retrieved from the Genome Warehouse (accession: GWHBCLC00000000; accessed November 28, 2023) following **Zhang et al. (2021)**.

Laccase (LAC) protein sequences from *O. sativa* (Oryzabase), *Arabidopsis thaliana* (TAIR), and *Theobroma cacao* (Phytozome) were used as references. A total of 74 LAC sequences were used to construct a Hidden Markov Model (HMM) with HMMER v3.2, which was then used to query the jute protein dataset to identify putative *C. capsularis* laccase (*CcaLAC*) genes.

Candidate sequences were validated for the presence of conserved laccase domains (PF07731.11, PF00394.19, PF07732.12) using NCBI CDD, HMMScan, and SMART.

Redundant hits (cut-off > 0.01) were removed to retain non-redundant sequences. Physicochemical properties were predicted using ExPASy ProtParam, and transmembrane regions were analysed using TMHMM v2.0.

### 2.2. Chromosome Mapping, Phylogenetic and Motif Analysis

Chromosomal locations of *CcaLAC* genes were visualised using MapInspect v1.0 based on their genomic coordinates (accessed November 28, 2023). Phylogenetic analysis was performed using laccase protein sequences from *C. capsularis*, *O. sativa*, *A. thaliana*, and *T. cacao*. *Arabidopsis* was included as a model dicot, rice as a representative monocot, and cacao because of its close genomic similarity to jute (**Zhang et al., 2021**). Multiple sequence alignment was performed using MUSCLE, and phylogenetic trees were constructed in MEGA v10.0 using the maximum-likelihood method with 1000 bootstrap replicates, gap deletion, and a site-gamma (G) distribution model. Conserved motifs in CcaLAC proteins were identified using the MEME suite, with parameters set to a maximum of 10 motifs, minimum motif width of 6, and maximum width of 50 (**Chanwala et al., 2023**).

### 2.3. Gene Structure, Duplication, and Synteny Analysis

The exon–intron organization of *CcaLAC* genes was analysed using the Gene Structure Display Server (GSDS). Gene duplication and mutation events within the *CcaLAC* family were investigated using BLASTp against laccase proteins from *A. thaliana, O. sativa, T. cacao,* and *C. capsularis*, with an e-value cutoff of 0.01. Gene duplication events were defined based on ≥80% sequence similarity (**Jha et al., 2021**). To assess evolutionary divergence, synonymous (Ks) and nonsynonymous (Ka) substitution rates were calculated using the TBtools program (**Chen et al., 2020**). Syntenic relationships among laccase genes were identified and visualised using TBtools.

### 2.4. Protein–Protein Interaction and Structural Prediction

Protein–protein interaction (PPI) networks for the candidate laccase proteins CcaLAC28 and CcaLAC32 were predicted using the STRING database (accessed April 2, 2025). Their *A. thaliana* orthologs, AtLAC4 and AtLAC17, respectively, were used as references for interaction inference. Secondary and tertiary structures of CcaLAC28 and CcaLAC32 were predicted using the Phyre2 server (accessed April 1, 2025). The predicted three-dimensional models were visualised and analysed using ChimeraX software.

### 2.5. Cis-Element and microRNA Target Analysis

Upstream promoter regions (∼2000 bp) of *CcaLAC* genes were retrieved from the *C. capsularis* genome for cis-regulatory element analysis. Putative cis-elements were identified using the PlantCARE database, with emphasis on elements associated with biotic and abiotic stress responses, growth and development, cell wall biosynthesis, circadian regulation, metabolism, and hormone signalling.

Arabidopsis mature microRNA (miRNA) sequences were obtained from the PmiREN2.0 database. Target sites of these miRNAs on *CcaLAC* genes were predicted using the psRNATarget server.

### 2.6. Gene Ontology Annotation

Gene Ontology (GO) annotation was performed to infer the functional roles of the *CcaLAC* gene family. *CcaLAC* protein sequences were searched against the UniProt-SwissProt database using BLASTp to identify homologs. The corresponding UniProt IDs were submitted to the Gene Ontology database (accessed January 15, 2025) for GO term assignment, with *A. thaliana* laccase proteins used as reference annotations. The analysis was performed using default parameters set in the portal (https://geneontology.org, accessed January 15, 2025): Test type-Fisher’s Exact; Correction- Calculate False Discovery rate. Only results displayed for FDR *P* < 0.05 were considered for further analysis.

### 2.7. Plant Material and Abiotic Stress Treatments

Seeds of *C. capsularis* variety JRC321, together with the low-lignin *dlpf* mutant and its parental control variety JRC212, were obtained from the Central Research Institute for Jute and Allied Fibres (CRIJAF), India. JRC321 was used for tissue-specific expression profiling and abiotic stress treatments, while the JRC212/*dlpf* pair was used for developmental-stage and mutant-comparison expression analyses. Plants were grown under greenhouse conditions (30 ± 2 °C, ∼80% relative humidity, and a 13 h light/11 h dark photoperiod).

The *dlpf* mutant and its control JRC212 have been previously characterised in detail with respect to growth phenotype, fibre lignin/cellulose composition, and the enzymatic basis of the lignification deficiency (**Sengupta and Palit, 2004**); its transcriptomic profile has additionally been characterised by **Chakraborty et al. (2015)**, which forms the basis of the *in silico* expression analysis in Section 2.8.

To examine tissue-specific expression of *CcaLAC* genes, leaf, root, xylem, and phloem tissues were collected from JRC321 plants at 60 days after sowing (DAS). For developmental stage–specific expression analysis, phloem tissues from JRC212 and *dlpf* plants were harvested at 30, 60, 90, and 120 DAS.

Abiotic stress treatments were performed on 60-day-old JRC321 plants. Abscisic acid (ABA; 0.15 mM) and copper stress (0.10 mM CuSO ·H O) were applied following **Parida et al. (2024)**. Stem tissues were collected at 0, 2, 4, 8, 12, and 24 h post-treatment, using three biological replicates per time point. All samples were immediately frozen in liquid nitrogen and stored at −80 °C for RNA isolation.

### 2.8. *In Silico* Analysis of *CcaLAC* Gene Expression

RNA-seq short reads derived from bast (phloem) tissues of control JRC212 and *dlpf* mutant genotypes were retrieved from the NCBI Sequence Read Archive (accession numbers SRR1585372 and SRR1585373, respectively). Reads were mapped against the coding sequences (CDS) of the 32 identified *CcaLAC* genes.

Transcript abundance was estimated using the Trinity RNA-seq pipeline scripts *align_and_estimate_abundance.pl* and *abundance_estimates_to_matrix.pl* (accessed July 10, 2024). TMM-normalised read counts were log -transformed, and expression heatmaps were generated using MeV software (retrieved July 10, 2024).

### 2.9. Gene Expression Analysis by qRT-PCR

Total RNA was isolated from leaf, xylem, phloem, and root tissues of jute plants using a commercial RNA isolation kit (Macherey-Nagel, Germany). Approximately 100 mg of tissue was ground in liquid nitrogen, and RNA extraction was performed according to the manufacturer’s protocol, including on-column DNase treatment to remove genomic DNA contamination. RNA integrity was verified on a 2% agarose gel, and concentration and purity were measured using a NanoDrop 2000c spectrophotometer (Thermo Scientific, USA).

First-strand cDNA was synthesised from 1 μg of total RNA using a first-strand cDNA synthesis kit (Thermo Scientific, USA). Gene-specific primers were designed using PrimerQuest (Integrated DNA Technologies, USA) with default parameters (Tm 59–62 °C, GC content 35–65%, primer length 17–30 bp, and amplicon size 75–150 bp). Primer sequences are listed in **Supplementary Table 1**.

qRT-PCR was performed using a QuantStudio™ 5 Real-Time PCR system (Thermo Scientific, USA). The cDNA was diluted 1:9 before amplification. PCR conditions included an initial denaturation at 95 °C for 2 min, followed by 40 cycles of denaturation at 95 °C for 15 s, primer-specific annealing at 95 °C for 1 min, and extension at 95 °C for 15 s. Melt curve analysis was conducted from 60 °C to 95 °C to confirm amplification specificity. Jute ubiquitin-1 (*UBI*; GenBank accession GH985256) was used as the internal reference gene (**Parida et al., 2024**). Relative gene expression was calculated using the 2^−ΔΔCt^ method (**Livak and Schmittgen, 2001**). Three biological replicates were analysed, each with three technical replicates.

Eight lignification-related Arabidopsis laccase genes (*AtLAC2, AtLAC4, AtLAC5, AtLAC10, AtLAC11, AtLAC12, AtLAC15,* and *AtLAC17*) were selected based on previous reports (**Berthet et al., 2012**). Homologous *CcaLAC* genes were identified by BLAST analysis in TBtools, and genes with >60% sequence identity were selected for expression analysis (**Jha et al., 2024**). Expression of these homologous *CcaLAC* genes was evaluated in phloem and xylem tissues at 30, 60, 90, and 120 DAS to assess their role in lignification during jute development.

### 2.10. Statistical Analysis

C_t_ values were averaged across three technical replicates per biological replicate (*n* = 3) prior to calculating relative expression (2 ^ΔΔCt^) in Microsoft Excel. For each *CcaLAC* gene independently, expression levels were compared using one-way ANOVA (*F*-test, *P* ≤ 0.05) followed by Tukey’s multiple comparisons test in GraphPad Prism (v9.0) - across tissues (Figure 3), developmental stages (Figure 4), and stress time points (Figure 6), and separately at each developmental stage for the JRC212 vs. *dlpf* comparison (Figure 5B). Data are presented as mean ± standard error (SE). As each gene was treated as an independent targeted hypothesis test rather than part of an unbiased genome-wide screen, no correction for multiple comparisons was applied across the 32-gene set, consistent with standard practice for candidate-gene qRT-PCR panels.

## 3. Results

### 3.1. Gene Number, Structure, and Duplication Events in White Jute *Laccase* Genes (*CcaLAC*s)

A total of 34 putative *laccase* genes were initially identified in our analysis, unevenly distributed across the seven chromosomes of white jute and designated *CcaLAC1* to *CcaLAC34* based on their chromosomal locations. Later, in the analysis of conserved domains, two candidates, *CcaLAC19* and *CcaLAC29*, lacked a complete set of conserved cupredoxin domains characteristic of functional laccases. These genes were subsequently excluded from further analysis, leaving 32 genuine *CcaLAC* genes. Chromosome 7 contained the highest number of *CcaLAC* genes (nine genes), followed by chromosome 1 with seven genes and chromosome 2 with six genes. Chromosomes 4 and 6 each harboured three *CcaLAC* genes, whereas chromosomes 3 and 5 contained only two (**Figure** 1A).

**Figure 1.**
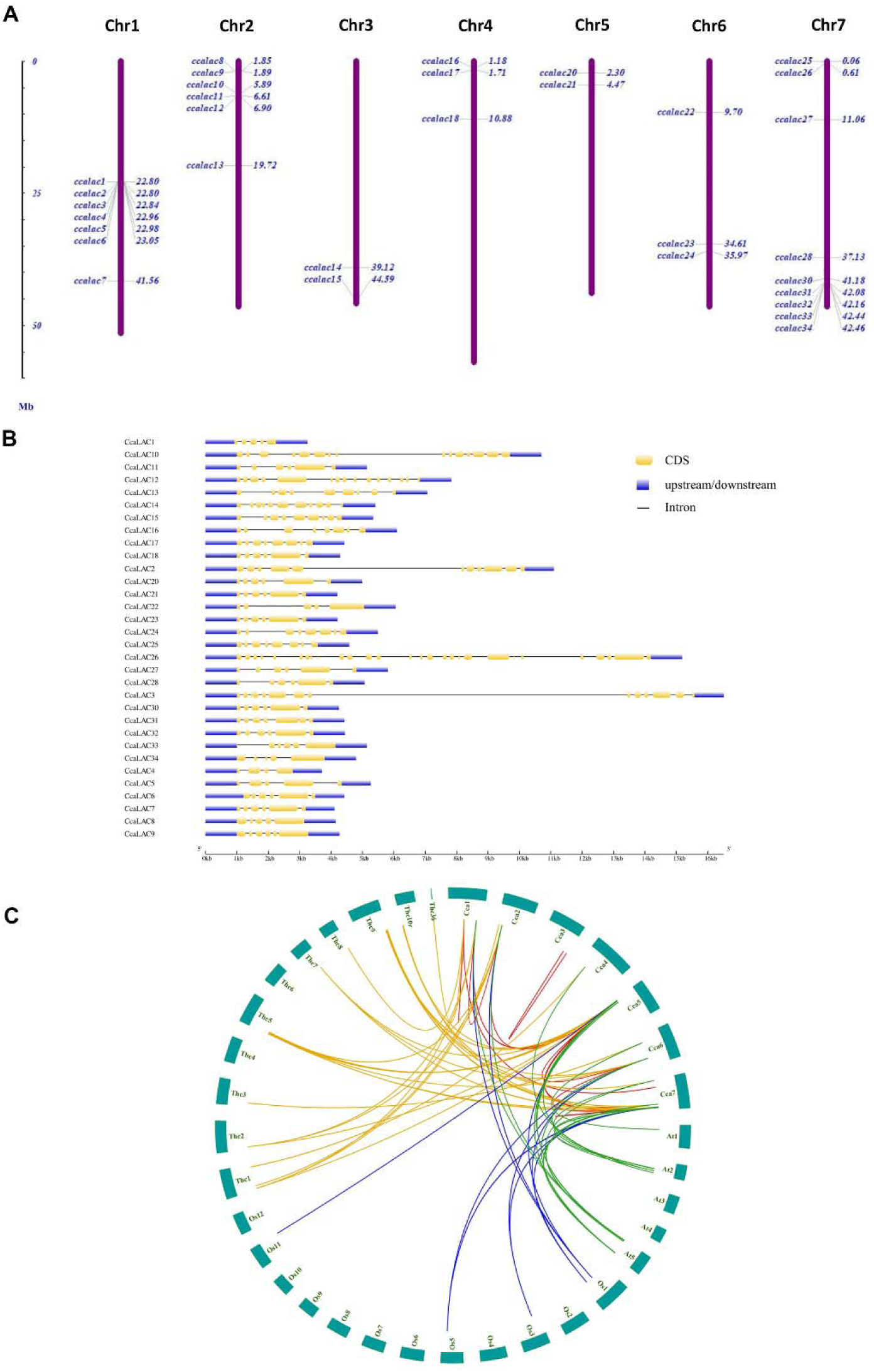
Number and gene structure of white jute (*C. capsularis*) laccase genes (*CcaLAC*s) (A) Chromosomal mapping of 32 laccase (*CcaLAC*) members in the genome. (B) Gene structure of CcaLACs. (C) Synteny relationship between white jute (*C. capsularis*), Arabidopsis (*A. thaliana*), rice (*O. sativa*), and cacao beans (*T. cacao*) *LAC* genes. Each coloured line represents duplicated gene pairs, indicating the shared ancestry among the species.

Substantial variation was observed in gene structure, with exon numbers ranging from four to 28 among the *CcaLAC* genes. *CcaLAC4* possessed the minimum number of exons (four), while *CcaLAC26* exhibited the most complex structure with 28 exons (**Figure** 1B).

Gene duplication analysis identified 18 paralogous *CcaLAC* gene pairs, including four tandem duplication events and fourteen segmental duplication events (**Supplementary Table 2**). In addition, 89 orthologous gene pairs were detected across species comparisons, comprising 18 pairs between *C. capsularis* and *O. sativa*, 20 pairs between *C. capsularis* and *A. thaliana*, and 51 pairs between *C. capsularis* and *T. cacao* (**Figure** 1C). The higher number of orthologous pairs between jute and cacao likely reflects their greater genomic similarity, as previously reported (**Zhang et al., 2021**). Ka/Ks analysis was successfully performed on the duplicated *CcaLAC* gene pairs, yielding reliable estimates. All analysed gene pairs exhibited Ka/Ks values <1, indicating that the duplicated genes have evolved under purifying selection (**Supplementary Table 5**).

### 3.2. Motif Analysis and Phylogenetic Relationships of Laccase Proteins in White Jute

The CcaLAC proteins exhibited considerable variation in size and physicochemical properties. Their molecular weights ranged from 54.01 kDa (CcaLAC16) to 177.31 kDa (CcaLAC26), with an average length of approximately 642 amino acids. The longest protein, CcaLAC26, consisted of 1,583 amino acids (**Table 1**). Most CcaLAC proteins displayed predicted instability indices below 40, indicating structural stability, except CcaLAC26 (43.34) (**Supplementary Table 4**). The aliphatic index ranged from 73.63 (CcaLAC27) to 93.14 (CcaLAC26), reflecting variability in thermal stability. All CcaLAC proteins showed negative average hydropathy values (<0), suggesting a predominantly hydrophilic nature. Subcellular localisation analysis predicted that CcaLAC proteins are distributed across multiple cellular compartments (**Supplementary Table 4**). Notably, 18 of the 32 CcaLAC proteins (56.25%) were predicted to possess transmembrane domains (**Supplementary Figure 1**).

**Table 1.** Bioinformatics Analysis and Physicochemical Properties of C*caLAC*s.

| Gene Name | Gene ID | Gene Length (bp) | Position | Amino Acid (aa) | pI | MW (kDa) | Subcellular Localization |
| --- | --- | --- | --- | --- | --- | --- | --- |
| *CcaLAC1* | GWHPBCLC001173 | 1330 | Chr1 | 293 | 5.01 | 31.85 | Chloroplast |
| *CcaLAC2* | GWHPBCLC001174 | 9172 | Chr1 | 1020 | 6.03 | 113.52 | Chloroplast |
| *CcaLAC3* | GWHPBCLC001176 | 14584 | Chr1 | 1007 | 5.07 | 111.63 | Endoplasmic Reticulum |
| *CcaLAC4* | GWHPBCLC001178 | 1789 | Chr1 | 381 | 8.18 | 42.81 | Cytoplasm |
| *CcaLAC5* | GWHPBCLC001179 | 3342 | Chr1 | 566 | 7.31 | 63.59 | Chloroplast |
| *CcaLAC6* | GWHPBCLC001182 | 2425 | Chr1 | 603 | 8.64 | 67.82 | Chloroplast |
| *CcaLAC7* | GWHPBCLC002591 | 2192 | Chr1 | 563 | 8.45 | 62.29 | Cytoplasm |
| *CcaLAC8* | GWHPBCLC003786 | 2149 | Chr2 | 588 | 8.69 | 66.19 | Mitochondria |
| *CcaLAC9* | GWHPBCLC003791 | 2275 | Chr2 | 611 | 6.55 | 68.30 | Cytoplasm |
| *CcaLAC10* | GWHPBCLC004194 | 8705 | Chr2 | 1027 | 9.54 | 114.71 | Cytoplasm |
| *CcaLAC11* | GWHPBCLC004256 | 3145 | Chr2 | 580 | 7.27 | 64.94 | Vacuoles |
| *CcaLAC12* | GWHPBCLC004295 | 5830 | Chr2 | 862 | 8.85 | 95.54 | Cytoplasm; Nucleus |
| *CcaLAC13* | GWHPBCLC005147 | 5065 | Chr2 | 594 | 6.07 | 66.39 | Vacuoles |
| *CcaLAC14* | GWHPBCLC008300 | 3416 | Chr3 | 635 | 8.72 | 71.13 | Golgi apparatus; Plasma Membrane |
| *CcaLAC15* | GWHPBCLC008874 | 3344 | Chr3 | 622 | 6.92 | 69.90 | Extracellular Space |
| *CcaLAC16* | GWHPBCLC009134 | 4102 | Chr4 | 485 | 9.04 | 54.01 | Golgi Apparatus |
| *CcaLAC17* | GWHPBCLC009197 | 2419 | Chr4 | 555 | 9.12 | 62.23 | Golgi Apparatus |
| *CcaLAC18* | GWHPBCLC010239 | 2289 | Chr4 | 575 | 8.39 | 63.76 | Extracellular Space |
| *CcaLAC20* | GWHPBCLC013482 | 2995 | Chr5 | 576 | 9.3 | 63.87 | Peroxisome |
| *CcaLAC21* | GWHPBCLC013679 | 2207 | Chr5 | 561 | 9.37 | 62.58 | Cytoplasm |
| *CcaLAC22* | GWHPBCLC016373 | 4059 | Chr6 | 578 | 6.83 | 64.63 | Cytoplasm |
| *CcaLAC23* | GWHPBCLC017617 | 2210 | Chr6 | 577 | 8.86 | 63.68 | Extracellular Space |
| *CcaLAC24* | GWHPBCLC017798 | 3491 | Chr6 | 542 | 9.51 | 60.92 | Endoplasmic Reticulum |
| *CcaLAC25* | GWHPBCLC018843 | 2576 | Chr7 | 525 | 6.05 | 57.92 | Chloroplast |
| *CcaLAC26* | GWHPBCLC018907 | 13184 | Chr7 | 1583 | 6.66 | 177.31 | Nucleus |
| *CcaLAC27* | GWHPBCLC019742 | 3813 | Chr7 | 564 | 8.94 | 63.75 | Chloroplast |
| *CcaLAC28* | GWHPBCLC020872 | 3070 | Chr7 | 560 | 9.28 | 61.46 | Vacuoles |
| *CcaLAC30* | GWHPBCLC021267 | 2250 | Chr7 | 576 | 9.41 | 63.62 | Vacuoles |
| *CcaLAC31* | GWHPBCLC021382 | 2427 | Chr7 | 574 | 9.19 | 63.11 | Nucleus |
| *CcaLAC32* | GWHPBCLC021391 | 2443 | Chr7 | 584 | 9.33 | 64.46 | Nucleus |
| *CcaLAC33* | GWHPBCLC021419 | 3143 | Chr7 | 551 | 6.36 | 61.84 | Cytoskeletons |
| *CcaLAC34* | GWHPBCLC021420 | 2794 | Chr7 | 565 | 6.4 | 63.17 | Cytoplasm; Nucleus |

Motif analysis identified ten conserved motifs (motif-1 to motif-10) across the CcaLAC protein family (**Figure** 2A). Most motifs were conserved in the majority of CcaLAC proteins, although motif-3 and motif-6 were absent in several members. Repetitions of specific motifs were observed in CcaLAC2, CcaLAC3, and CcaLAC10, suggesting possible functional diversification.

**Figure 2.**
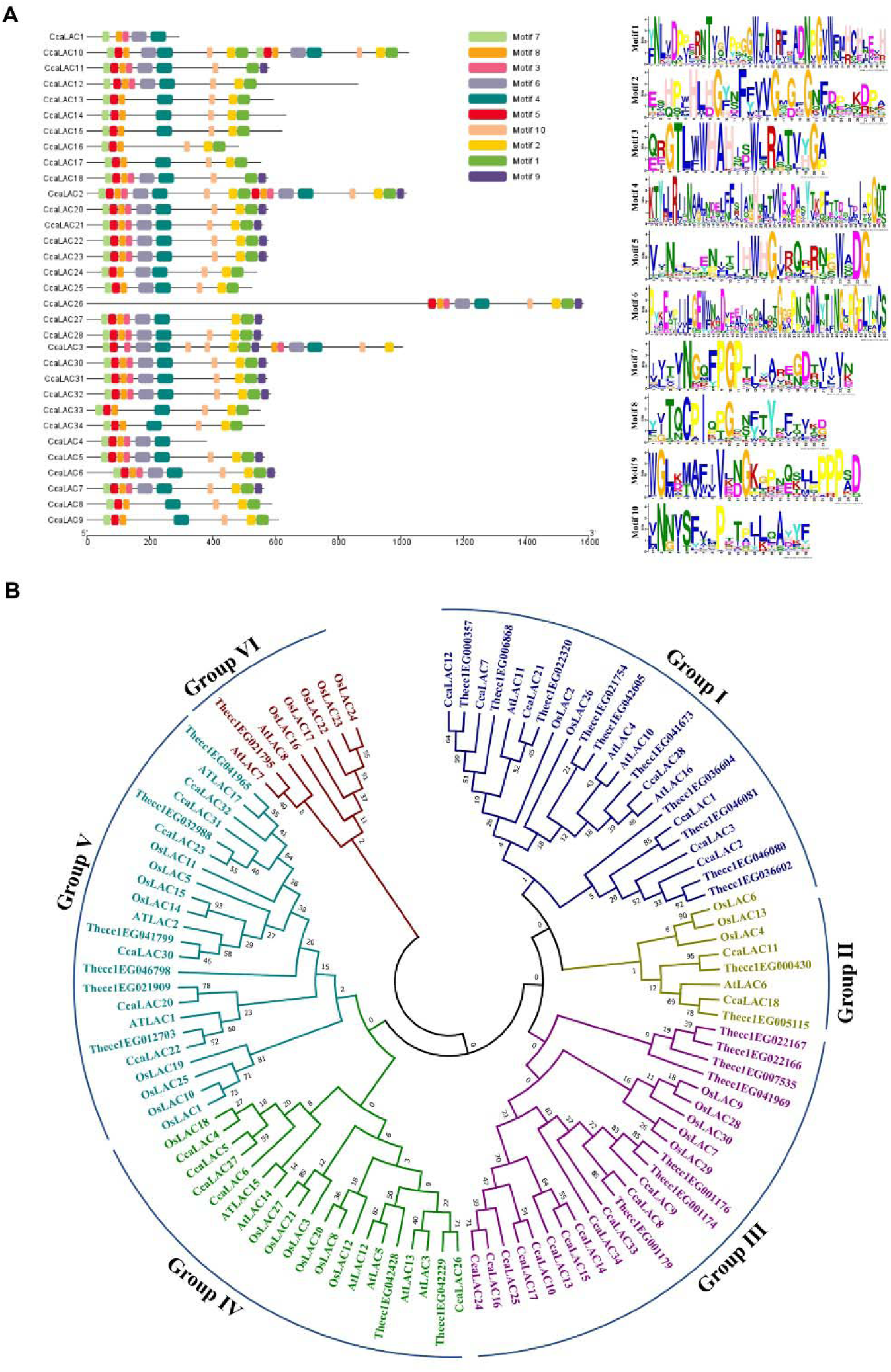
Conserved motif and phylogenetic analysis of CcaLAC proteins. (A) Conserved motif distribution in identified CcaLAC proteins. Different colours and their respective sequences indicate motifs. (B) Phylogenetic tree of LAC proteins of white jute (*C. capsularis*), rice (*Oryza sativa*), *A. thaliana*, and *T. cacao*. The analysis classified the members of the LAC protein family into six distinct groups. Abbreviations such as At (*Arabidopsis*), Os (rice), Thecc (*cacao*), and Cca (white jute) are used to denote laccase genes specific to each species.

Phylogenetic analysis grouped the 32 CcaLAC proteins into six distinct clades (Groups I–VI) (**Figure** 2B). Group III contained the largest number of members (12 proteins), followed by Group I (7 proteins), Group V (6 proteins), Group IV (5 proteins), and Group II (2 proteins). Group VI contained none of the CcaLAC protein. Functionally characterised Arabidopsis laccases clustered with their putative jute orthologs: AtLAC4, a key enzyme in lignification, grouped with CcaLAC28 in Group I, whereas AtLAC17 clustered with its jute homolog CcaLAC32 in Group V (**Figure** 2B). The **Supplementary Table 3** contain information on bootstrap support values for phylogenetic clades, their descendant taxa, and phylogenetic groupings.

### 3.3. Cis-acting Elements

A diverse array of cis-acting regulatory elements was identified in the upstream promoter regions of the 32 *CcaLAC* genes (**Supplementary Figure 2**). Core promoter elements such as MYB, MYC, and TATA-box motifs were present in all *CcaLAC* promoters. In addition, cis-elements associated with stress responsiveness (e.g., STRE, W-box, MYB, MYC, and WUN-motif), hormonal signalling (e.g., ABRE, ERE, P-box, GARE-motif, and TGA element), and tissue-specific and developmental regulation (e.g., G-box, GT1-motif, RY-element, Sp1, GATA-motif, LAMP element, Box-4, and TCT-motif) were abundant, suggesting functional diversification and complex regulatory control of *CcaLAC* gene expression (**Ain-Ali et al., 2021**; **Jha et al., 2024**).

Notably, several cis-elements implicated in secondary cell wall formation were detected upstream of multiple *CcaLAC* genes (**Supplementary Table 6**). These included AC elements (ACE), MYB-binding sites (MBS), and G-box motifs, which are known to regulate lignin biosynthesis and secondary wall development, as well as to play roles in hormonal signalling (**Zhou et al., 2009**). The presence of these elements supports a potential involvement of *CcaLAC* genes in cell wall formation and lignification processes in white jute.

### 3.4. Tissue-Specific Expression Analysis of *CcaLAC* Genes

qRT-PCR analysis revealed detectable expression of all 32 *CcaLAC* genes across four tissues—phloem, xylem, leaf, and root. Transcript abundance was quantified as relative expression (RQ) values, with root tissue expression set as the reference (RQ = 1) (**Figure** 3).

**Figure 3.**
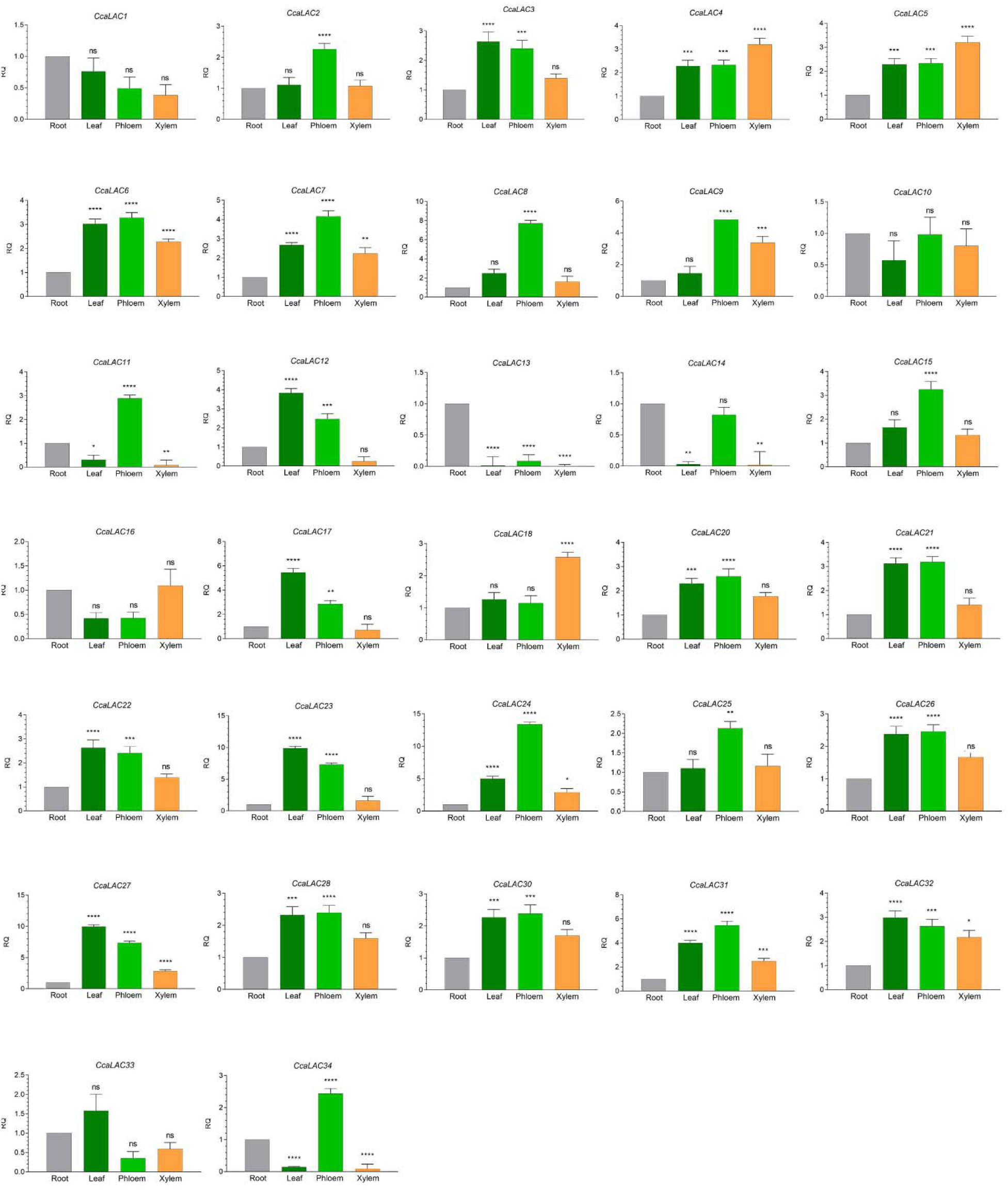
Tissue-specific expression of laccase (*CcaLAC*) genes. This figure shows the relative quantification (RQ) values for the *CcaLAC* gene expression in the leaf, phloem, xylem, and root tissues. The RQ values are normalised using the ubiquitin 1 (*UBI*) housekeeping gene. A one-way ANOVA (F-test; *P* ≤ 0.05) was performed independently for each gene, followed by Tukey’s multiple comparisons test. Asterisks denote significant differences (*P* ≤ 0.05); ’ns’ indicates non-significant differences. Data represent mean ± SE of three biological replicates. No correction for multiple comparisons was applied across genes.

Several *CcaLAC* genes exhibited higher expression in leaf tissue, including *CcaLAC3, CcaLAC12, CcaLAC17, CcaLAC22, CcaLAC23, CcaLAC27, CcaLAC32,* and *CcaLAC33*.

Enhanced expression in phloem tissue was observed for *CcaLAC2, CcaLAC6, CcaLAC7, CcaLAC8, CcaLAC9, CcaLAC11, CcaLAC15, CcaLAC20, CcaLAC21, CcaLAC24, CcaLAC25,*

*CcaLAC26, CcaLAC28, CcaLAC30, CcaLAC31,* and *CcaLAC34*. Higher transcript accumulation in xylem tissue was recorded for *CcaLAC4, CcaLAC5, CcaLAC16,* and *CcaLAC18*, whereas *CcaLAC1, CcaLAC10, CcaLAC13,* and *CcaLAC14* showed relatively higher expression in root tissue.

### 3.5. Phloem Tissue-Specific Expression Analysis of *CcaLAC* Genes Across Developmental Stages

Expression profiling was performed at four key time points: 30 DAS, corresponding to the initiation of fibre development, and 60, 90, and 120 DAS, which represent progressive stages leading to fibre maturation and harvest. Several *CcaLAC* genes, including *CcaLAC21, CcaLAC23, CcaLAC28,* and *CcaLAC32*, exhibited a steady increase in transcript abundance from 30 DAS to 120 DAS (**Figure** 4). In contrast, *CcaLAC12, CcaLAC20, CcaLAC26,* and *CcaLAC30* showed a gradual increase in expression up to 90 DAS, followed by a marked decline at 120 DAS. Meanwhile, *CcaLAC7* and *CcaLAC31* displayed a slight reduction in expression at 120 DAS compared to earlier stages.

**Figure 4.**
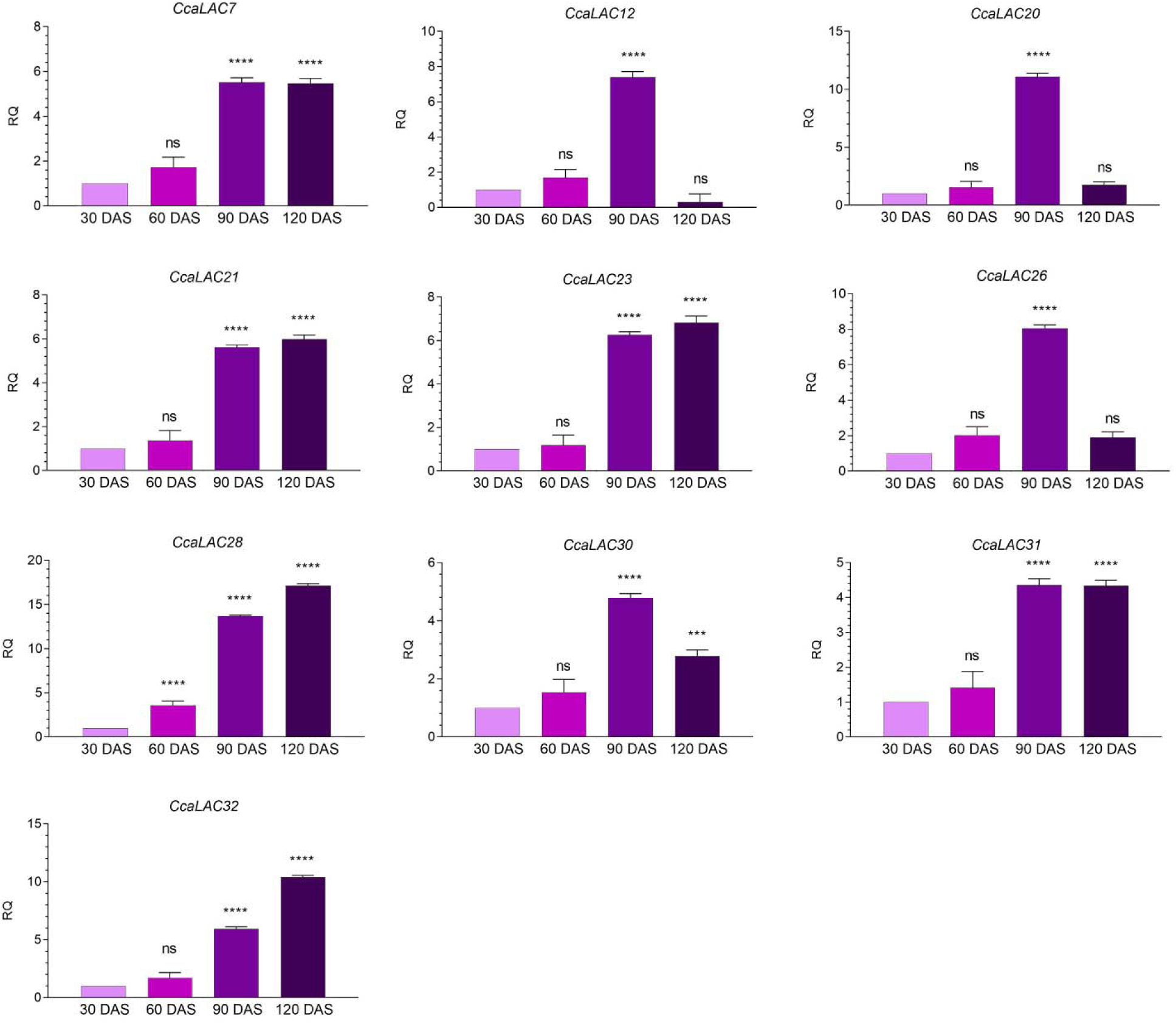
Expression of laccase (*CcaLAC*) genes at different developmental stages of white jute plants. This figure illustrates the expression of *CcaLAC*s in the phloem tissue of white jute at four different time points, from 30 to 120 days after sowing (DAS). A one-way ANOVA (F-test; *P* ≤ 0.05) was performed independently for each gene, followed by Tukey’s multiple comparisons test. Asterisks denote significant differences (*P* ≤ 0.05); ’ns’ indicates non-significant differences. Data represent mean ± SE of three biological replicates. No correction for multiple comparisons was applied across genes.

Overall, the majority of the analysed *AtLAC* homologous genes associated with the lignin biosynthetic pathway demonstrated increased expression in phloem tissue from 30 DAS to 90 DAS, coinciding with the active phase of fibre development and lignification.

### 3.6. Expression Analysis of *CcaLAC* Genes in the Low-Lignin Mutant Jute (*dlpf*)

*In silico* expression analysis of *CcaLAC* genes was performed using RNA-seq data from 30 DAS phloem tissue of the white jute cultivar JRC212 and its X-ray-irradiated low-lignin mutant, *dlpf*, sourced from **Chakraborty et al. (2015)**. All 32 *CcaLAC* genes were expressed in phloem tissue (**Figure 5A**). Compared with JRC212, *dlpf* showed reduced expression of *CcaLAC3, CcaLAC4, CcaLAC7, CcaLAC9, CcaLAC17, CcaLAC18, CcaLAC21, CcaLAC23, CcaLAC31, CcaLAC32,* and *CcaLAC33*. To validate the RNA-seq results, eight *CcaLAC* genes (*CcaLAC3, CcaLAC4, CcaLAC7, CcaLAC11, CcaLAC17, CcaLAC21, CcaLAC31,* and *CcaLAC32*) were randomly selected for qRT-PCR analysis, which confirmed expression patterns consistent with the transcriptomic heatmap (**Supplementary Figure 3**).

**Figure 5.**
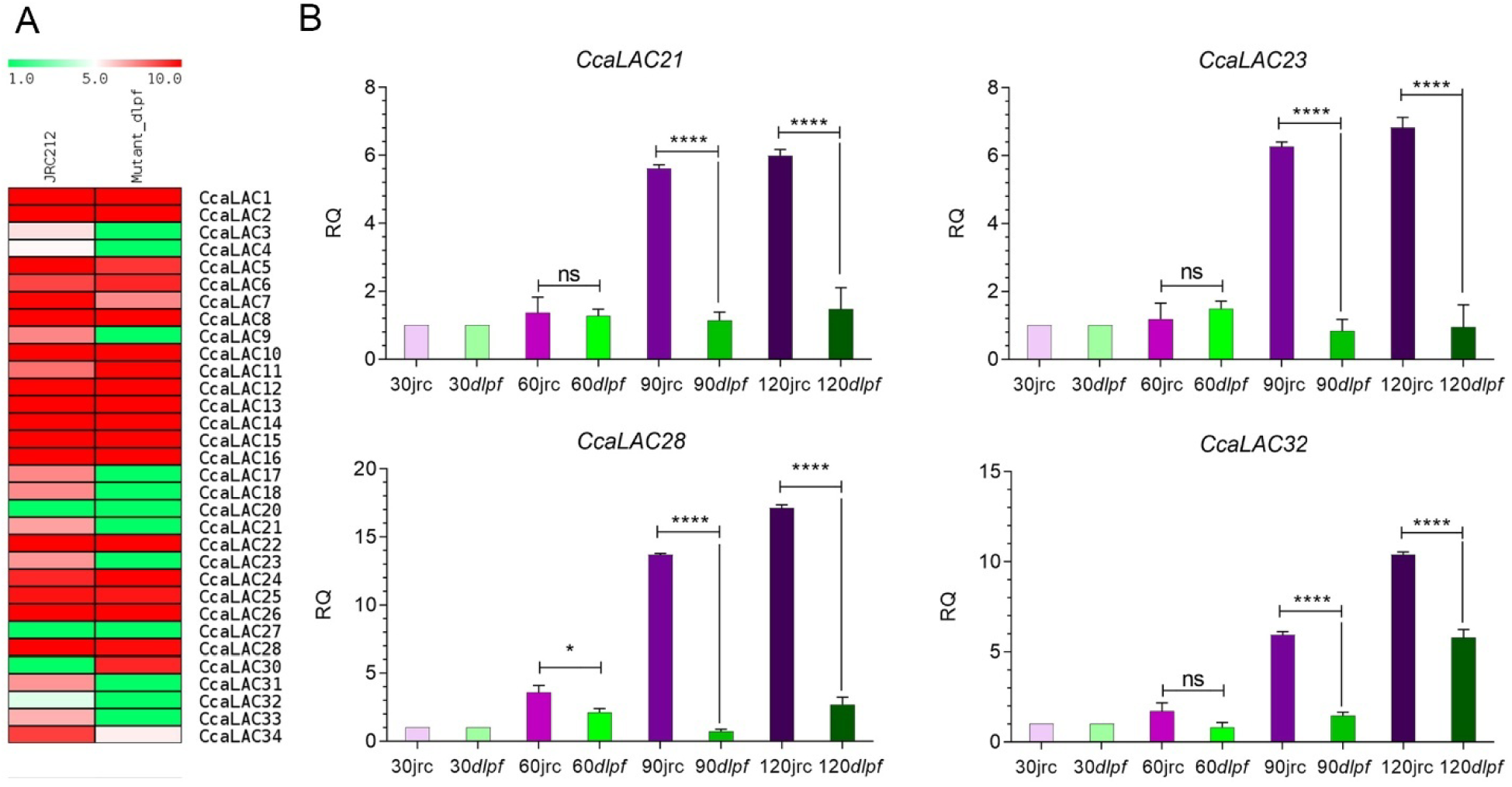
Expression analysis of white jute *laccase* genes in phloem tissue. (A) Expression analysis *in silico* of *CcaLAC*s in the phloem tissue of JRC212 (control) and *dlpf* (mutant) at 30 DAS. The heat map is based on publicly available transcriptomics data from BioProject PRJNA255934. The expression scale ranges from low (1, green) to high (10, red). (B) This figure presents a comparative expression analysis of laccase (*CcaLAC*) genes of JRC212 and *dlpf* (which has fibres with 50% lower lignin content), across different developmental stages (30 DAYS to 120 DAS). A separate one-way ANOVA (F-test; *P* ≤ 0.05) comparing JRC212 and *dlpf* was performed independently at each developmental stage (30, 60, 90, and 120 DAS), followed by Tukey’s multiple comparisons test. Asterisks denote significant differences (*P* ≤ 0.05); ’ns’ indicates non-significant differences. Data represent mean ± SE of three biological replicates.

Developmental expression analysis revealed that *CcaLAC21, CcaLAC23, CcaLAC28*, and *CcaLAC32*—genes that showed a progressive increase in expression from 30 to 120 DAS in JRC212—were significantly downregulated in *dlpf* at 90 and 120 DAS (*P* < 0.0001; **Figure** 5B). Similarly, *CcaLAC12, CcaLAC20,* and *CcaLAC26*, which exhibited peak expression at 90 DAS in JRC212, showed reduced expression in *dlpf* at the same stage (**Supplementary Figure 4**).

### 3.7. Expression Analysis of *CcaLAC* Genes Under Copper Stress and ABA Treatment

*CcaLAC28* and *CcaLAC32*, homologous to *A. thaliana AtLAC4* and *AtLAC17*, respectively, are among the highly expressed *CcaLAC* genes and are implicated in lignin biosynthesis. To evaluate their stress responsiveness, expression profiles were analysed in response to copper and abscisic acid (ABA) treatments.

Under copper stress, *CcaLAC28* showed a significant increase in expression at 12 h, whereas *CcaLAC32* exhibited significantly elevated expression at all examined time points (2–24 h). *CcaLAC32* expression increased gradually, peaked at 8 h, and subsequently declined (**Figure** 6A).

**Figure 6.**
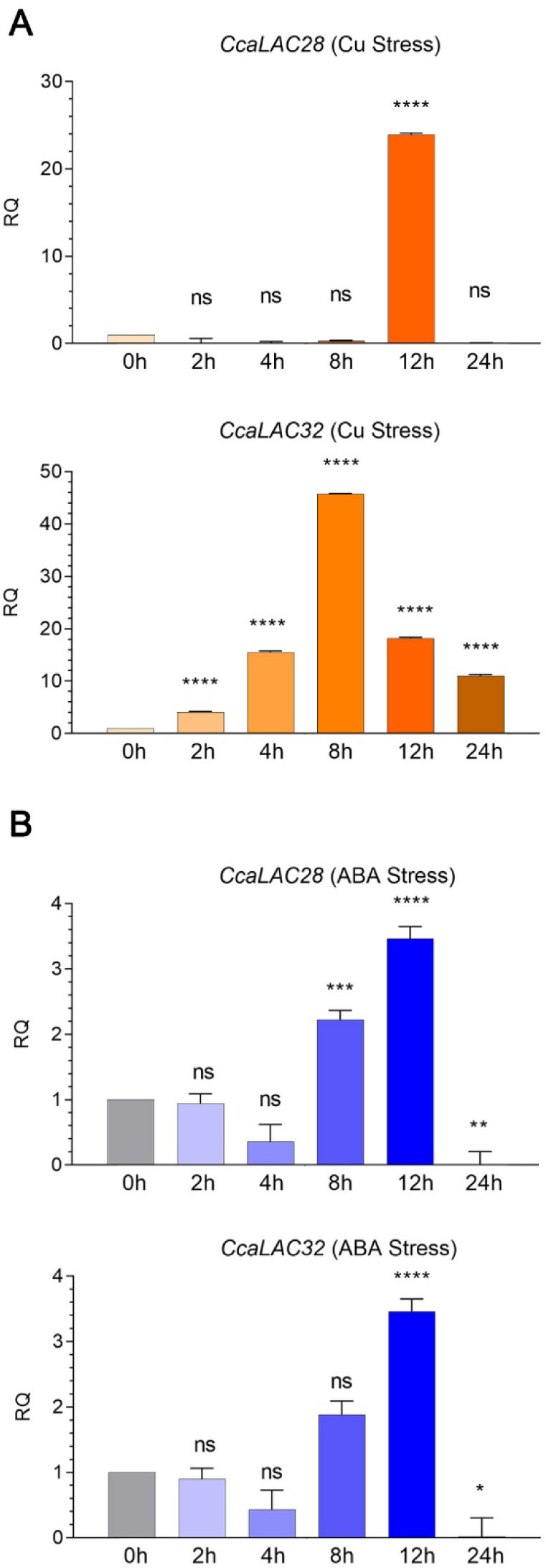
Expression of *CcaLAC28* and *CcaLAC32* under abiotic stresses. This figure depicts the *CcaLAC28* and *CcaLAC32* gene expression levels under (A) copper stress (0.10 mM CuSO_4_·H_2_O) and (B) ABA hormone stress. A one-way ANOVA (F-test; *P* ≤ 0.05) was performed independently for each gene, followed by Tukey’s multiple comparisons test. Asterisks denote significant differences (*P* ≤ 0.05); ’ns’ indicates non-significant differences. Data represent mean ± SE of three biological replicates. No correction for multiple comparisons was applied across genes.

Under ABA treatment, both *CcaLAC28* and *CcaLAC32* showed significant induction at 8 h and 12 h, with maximum expression at 12 h, followed by a decline at 24 h (**Figure** 6B). These expression patterns suggest that *CcaLAC28* and *CcaLAC32* function as delayed-responsive genes in white jute under ABA-mediated stress.

### 3.8. Protein Structure Analysis of CcaLAC28 and CcaLAC32

The protein structures of CcaLAC28 and CcaLAC32 were predicted using Phyre2.2. CcaLAC28 (531 amino acids) was modelled with 100% confidence and showed 48% sequence identity with the crystal structure of *Z. mays* laccase 3 (ZmLAC3) complexed with sinapyl (fold library ID: c6kliA). The predicted secondary structure consisted of 39% β-strands, 4% α-helices, and 7% disordered regions, with a transmembrane domain predicted between residues 226 and 241 (**Figure** 7A). Similarly, CcaLAC32 (540 amino acids) was modelled with 100% confidence and exhibited 50% sequence identity with ZmLAC3. Its predicted secondary structure comprised 34% β-strands, 5% α-helices, and 4% disordered regions, with a transmembrane domain predicted between residues 136 and 151 (**Figure** 7B).

**Figure 7.**
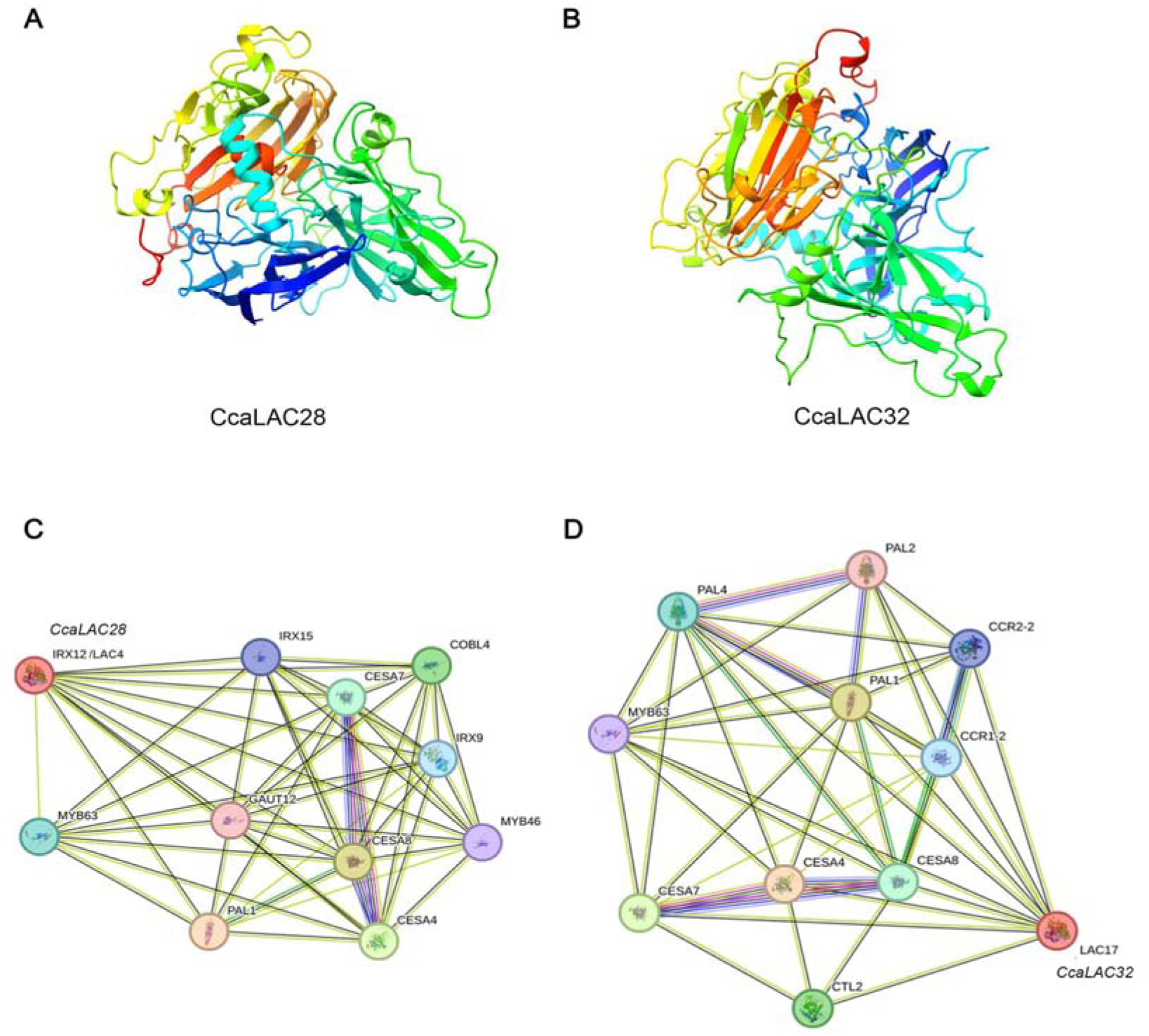
Predicted three-dimensional protein structures and Protein-protein interaction network analysis of CcaLAC28 and CcaLAC32. (A) The structural model of CcaLAC28 (531 amino acids) and (B) CcaLAC32 (540 amino acids) were generated using Phyre2.2. The structures are colour-coded from the N-terminus (blue) to the C-terminus (red), highlighting secondary structural elements such as β-strands, α-helices, and loop regions. (C) Predicted interaction network of *CcaLAC28* (homologous to *AtLAC4*). (D) Predicted interaction network of *CcaLAC32* (homologous to *AtLAC17*). The STRING database is used for protein-protein interaction network analysis.

### 3.9. Protein–Protein Interaction Network Analysis

Protein–protein interaction networks were predicted *in silico* using the STRING database. As the jute genome is not available in STRING, *Arabidopsis thaliana* (NCBI Taxonomy ID: 3702) was used as a reference to analyse the interaction networks of AtLAC4 and AtLAC17, homologous to CcaLAC28 and CcaLAC32, respectively.

CcaLAC28/AtLAC4 was predicted to interact with several enzymes and regulators associated with lignin biosynthesis and secondary cell wall formation, including phenylalanine ammonia-lyase (PAL1), cellulose synthase catalytic subunits (CESA4, CESA7, and CESA8), and transcription factors MYB63 and MYB46. Additional interactions were observed with proteins involved in secondary wall assembly, such as IRX9, IRX15, COBRA-like protein 4 (COBL4), and galacturonosyltransferase 1 (GAUT1) (**Figure** 7C).

Similarly, CcaLAC32/AtLAC17 was predicted to interact with PAL1, PAL2, PAL4, CESA4, CESA7, CESA8, and MYB63. In addition, unique interactions were identified with cinnamoyl-CoA reductase 1 (CCR1-2), CCR2-2, and chitinase-like protein 2 (CTL2), indicating a potential role in lignin biosynthesis, stress response, and cell wall organisation (**Figure** 7D).

### 3.10. MicroRNA (miRNA) Targeting of *CcaLAC* Genes

Several *CcaLAC* genes were predicted as targets of the lignin-associated miRNA miR397 (**Supplementary Table 7; Supplementary Figure 5**). Ath-miR397a was predicted to target *CcaLAC2, CcaLAC3, CcaLAC7, CcaLAC20, CcaLAC21, CcaLAC22, CcaLAC23, CcaLAC26, CcaLAC28, CcaLAC30,* and *CcaLAC31*, whereas Ath-miR397b targeted *CcaLAC7, CcaLAC20, CcaLAC21, CcaLAC22, CcaLAC23, CcaLAC28, CcaLAC30, CcaLAC31,* and *CcaLAC34*.

### 3.11. Functional Annotation of *CcaLAC* Genes by Gene Ontology (GO) Analysis

GO term analysis indicated that *CcaLAC* genes are primarily involved in lignin biosynthesis and metabolism, including lignin biosynthetic (GO:0009809), lignin metabolic (GO:0009808), phenylpropanoid biosynthetic (GO:0009699), and phenylpropanoid metabolic processes (GO:0009698) (**Supplementary Figure 6**). Additional enrichment was observed for secondary metabolite biosynthesis and metabolism. At the molecular function level, *CcaLACs* were associated with oxidoreductase activity, copper ion binding, and transition metal ion binding, consistent with the conserved biochemical roles of plant laccases (**Supplementary Figure 6**).

## 4. Discussion

Beyond extending genome-wide *LAC* characterisation in white jute (*C. capsularis*), this study is the first to directly link *CcaLAC* expression to a lignin-deficient phenotype (*dlpf*) and to abiotic stress responsiveness, thereby allowing candidate genes to be prioritised on convergent rather than purely homology-based evidence. The white jute genome contains 25,874 genes (**Zhang et al., 2021**), including the largely unexplored laccase (*CcaLAC*) genes, particularly regarding their roles in stem development, fibre lignification, and performance under abiotic and biotic stress. The research aimed to determine the functional importance of *LAC* in white jute. The entire white jute genome was searched, and 32 *CcaLAC* genes were identified; their expression was examined across different tissues (**Figure 3**). Out of the 32 *CcaLAC* genes, 16 were highly expressed in phloem, 8 in leaves, 4 in xylem, and 4 in roots. Considering the stem part as a combination of phloem and xylem tissue-specific expression, 61.76% of the *CcaLAC* genes are expressed here, which is highly significant in terms of the role of laccase in jute stem and fibre development. Such observations have been reported in which *AtLAC* genes (*AtLAC2, AtLAC4, AtLAC10, AtLAC11, AtLAC16,* and *AtLAC17*), responsible for lignification, were overexpressed in Arabidopsis stem tissue (**Berthet et al., 2012**). This tissue-specific expression pattern of *LAC* genes in white jute (*C. capsularis*) differs from that observed in tossa jute (*C. olitorius*). In tossa jute, 43.47% (20 of 46) of the *ColLAC* genes showed higher expression in roots than in other tissues tested (**Jha et al., 2025**). This suggests potential differences in regulatory mechanisms between the two *Corchorus* species that govern the tissue-specific expression patterns of *LAC* genes.

Separation of xylem and phloem tissues from the jute stem is difficult during the initial days of growth, but by 30 DAS it becomes much easier, suggesting that fibre development and lignification begin around this time (**Chakraborty et al., 2015**). By 110-120 DAS, phloem fibre can be easily extracted via biological retting, indicating its maturity and the completion of lignification. This research targeted monthly time points of 30 DAS (one month), 60 DAS (two months), 90 DAS (three months), and finally 120 DAS (four months) to relate *CcaLAC* expression to jute stem development and the fibre lignification process.

As a part of this investigation to find the functional importance of *CcaLAC* in the fibre lignification process of jute, homologous genes from *Arabidopsis* were selected. *Arabidopsis* is commonly used for lignin pathway analysis in plants due to its lignified stem tissue, short life span, and its classification as a dicotyledonous plant (**Vanholme et al., 2010**). There is a remarkable similarity between the sequences of *CcaLAC* and *AtLAC* genes, and their expression patterns also match those of *Arabidopsis*, increasing with plant development (**Berthet et al., 2011**) (**Figure 4**).

Mutation of Arabidopsis *AtLAC4* (homologous to *CcaLAC28*) and *AtLAC17* (homologous to *CcaLAC32*) individually resulted in a 10% reduction in lignin content, and when both genes (*AtLAC4+AtLAC17*) were mutated, lignin content was reduced by 39% (**Berthet et al., 2011**). This same study on the double mutant line also demonstrated that *AtLAC4* and *AtLAC17* contributed significantly to total laccase activity in plants, with a ∼50% reduction in laccase activity upon their mutation. Recently, overexpression of *AtLAC4* in populus increased lignin content in the transgenic populus without any phenotypic changes in the plants (**Ahlawat et al., 2024**). These findings established the involvement of *AtLAC* in the lignification process, specifically *AtLAC4* and *AtLAC17*. This suggests that their homologous genes, *CcaLAC28* and *CcaLAC32*, might also contribute significantly to lignification. The transcriptional expression of *CcaLAC* genes was assessed at different developmental stages (30 DAS to 120 DAS) in jute plants using qRT-PCR to quantify mRNA transcripts. Four genes—*CcaLAC21, CcaLAC23, CcaLAC28,* and *CcaLAC32*—showed a steady increase in expression as the plant developed (**Figure 4**). This raised the question: Are these genes important in lignin production in jute? Finding the answer led us to examine the expression of these four *CcaLAC* genes in a low-lignin-fibre-content mutant variety of white jute, *dlpf,* at various stages of growth. We found statistically significant differences in the expression patterns of these four CcaLAC genes between the low-lignin mutant, *dlpf*, and its control, JRC212. Lower expression patterns of *CcaLAC21, CcaLAC23, CcaLAC28* and *CcaLAC32* were found, which could be a contributing factor to the low lignin concentration observed in *dlpf* (**Figure 5B**). Although *dlpf* is an X-ray-irradiated mutant and may have mutations in other lignin pathway genes, the differential expression pattern of its control (JRC212) on four *CcaLAC* genes (*CcaLAC21, CcaLAC23, CcaLAC28,* and *CcaLAC32*) suggests that these genes are potential candidates for further investigation.

The expression of *CcaLAC12, CcaLAC20,* and *CcaLAC26* was significantly higher at 90 DAS, but decreased at 120 DAS to levels similar to those observed at 30 DAS (**Supplementary Figure 4**). If these genes were solely involved in the lignification process, their expression should remain high until 120 DAS (as in *CcaLAC21, CcaLAC23, CcaLAC28,* and *CcaLAC32*). The observed decrease in expression suggests a potential role in stem elongation or structure, which 90 DAS might achieve. Further functional characterisation is necessary to confirm this hypothesis. This expression pattern was also observed in the *dlpf* mutant (**Supplementary Figure 4**).

Previous studies (up to 60 DAS) on the low-lignin mutant jute, *dlpf*, by **Chakraborty et al. (2015)** reported low expression of cinnamyl alcohol dehydrogenase (CAD), which is considered essential for the lignin pathway. This recent study on the *dlpf* (plants up to 120 DAS) lignin pathway involved comparing the qRT-PCR expression data using two housekeeping genes- *UBI* and *beta-tubulin* (*BTUB*) (**Parida et al., 2024)**. We found the expression of *CCoAoMT*, cinnamate 4-hydroxylase (*C4H*), and *CAD* to be lower in *dlpf* than in its control JRC212. This study represents the first analysis of *LAC* gene expression in the low-lignin mutant *dlpf*. Although transcriptomic data (BioProject PRJNA255934) from **Chakraborty et al. (2015)** reported 15 unigenes annotated as *LAC* genes in *JRC212* and *dlpf*, no qRT-PCR-based expression analysis was conducted. Here, we utilised the same transcriptomic dataset to examine the *in silico* expression profiles of all 32 *CcaLAC* genes, revealing distinct expression patterns (**Figure 5A**). The low expression levels of *CcaLAC21, CcaLAC23, CcaLAC28* and *CcaLAC32* in *dlpf* may also contribute to the reduced lignin content (**Figure 5B**). As statistical comparisons were performed independently for each gene without correction across the 32-member *CcaLAC* family, the aggregate pattern of differentially expressed genes should be interpreted as indicative rather than as a formally corrected genome-wide result.

Consequently, the reduced expression of *CcaLAC21, CcaLAC23, CcaLAC28* and *CcaLAC32* observed in this mutant- *dlpf* cannot be interpreted as direct evidence of their causal role in lignin biosynthesis, as the altered expression may result from other unidentified mutations or indirect regulatory effects. However, the *dlpf* mutant has previously been transcriptomically characterised by **Chakraborty et al. (2015)**, who reported differential expression of several genes involved in secondary cell wall formation and lignin biosynthesis, including Phenylalanine ammonia-lyase (PAL), cinnamyl alcohol dehydrogenase (CAD), and multiple transcription factors. Building upon this well-characterised genetic resource, we reanalysed the publicly available RNA-seq dataset and found that *CcaLAC28* and *CcaLAC32* were also significantly downregulated, suggesting their potential association with lignification. Nevertheless, these findings should be considered correlative rather than causal. Given the limited availability of genetically defined lignin mutants in jute, this transcriptomically validated *dlpf* mutant provides a valuable resource for candidate gene discovery.

Transcriptional and post-transcriptional factors influence the expression pattern of plant laccases. The MYB transcription factors regulate *LAC*, as reported in multiple plant studies. They are among the largest transcription factor families and are highly functionally diverse. In Arabidopsis, *AtMYB58* directly regulates *AtLAC4* expression; *AtMYB46* regulates *AtLAC10,* and *AtHY5* regulates *AtLAC12* (**Zhou et al., 2009**). In bamboo (*Phyllostachys edulis*), *PeMYB4.1/20/85.2* regulates *PeLAC20*; in pear (*Pyrus bretschneideri*), *PbMYB26* regulates *PbLAC4*-like; and in Pearl millet (*Pennisetum glaucum*)*, PgMYB305* regulates *PgLAC14* (**Bai et al., 2023**). A predominance of MYB binding sites was found in many jute laccases (*CcaLAC* genes) after analysis of cis-elements in their upstream sequences (**Supplementary Figure 2**). MYBs were found upstream of *CcaLAC21, CcaLAC23, CcaLAC28,* and *CcaLAC32,* indicating that MYBs could regulate these genes. Recently, 88 MYBs were identified in white jute through a whole-genome search, but it remains to be determined which MYBs regulate *CcaLAC* expression (**Niyitanga et al., 2023**).

The role of miRNAs in regulating LAC has been reported in multiple plant species, including rice, Arabidopsis, and Populus. MicroRNA397 (miR397) directly targets laccase transcripts and regulates their expression in plants. In Populus, miR397 targets 28 *PtLAC* genes (**Lu et al., 2013**). In chickpea (*Cicer arietinum*), miR397 targets *CaLAC4* and *CaLAC17* among the 20 identified *CaLAC* genes (**Sharma et al., 2023**). Similarly, miR397 regulates *OsLAC* expression in rice (**Bakhshi et al., 2016**). It is considered a negative regulator of *LAC* expression in lignin biosynthesis, as reported in rice, chickpea, Arabidopsis, and Populus. Does miR397 target the jute *CcaLAC*s? *In silico* analysis predicted that miR397a targets 11 *CcaLAC*s, and miR397b targets 9 *CcaLAC*s, in white jute (**Supplementary Figure 5**). *CcaLAC28* was predicted to be targeted by both miR397a and miR397b, whereas no target site was predicted for *CcaLAC32* (homologous to *AtLAC17*). Overexpressing miR397 in jute could provide further insight into its involvement in the lignification process by regulating *CcaLACs* in the near future. These relationships are based on sequence-complementarity prediction using Arabidopsis miRNA sequences and have not been experimentally validated in jute; degradome sequencing or 5′-RACE analysis, together with confirmation of endogenous miR397 expression in jute tissues, will be required to establish genuine miR397–*CcaLAC* targeting relationships.

The assortment of cis-acting elements located in the upstream promoter regions of white jute *CcaLACs* suggests that the expression of *CcaLAC* genes may be influenced by diverse factors, including plant growth, developmental stages, hormonal signals, and both abiotic and biotic stresses (**Supplementary Figure 2**). Notably, cis-acting elements associated with secondary wall formation, such as AC elements (ACE), G-box motifs, and MYB Binding Sites (MBS), were identified in multiple *CcaLACs* (**Supplementary Table 6**). **Zhou et al. (2009)** demonstrated in *Arabidopsis* that *AtMYB58* and *AtMYB63* directly activate lignin biosynthetic genes by targeting ACE. **Zhao et al. (2010)** also reported that the lignin-specific transcription factor *AtMYB58* in *Arabidopsis* can directly activate most lignin biosynthetic genes through ACE binding in their promoters. Additionally, G-box motifs or G-box Regulating Factors (GRFs) play crucial roles in plant physiological and developmental processes, as well as in stress responses (**Sun et al., 2022**). The presence of these cis-acting elements in the *CcaLAC* members highlights their potential involvement in the lignification process and secondary wall formation in jute.

It is well-documented that laccase (*LAC*) genes are susceptible to environmental changes, with expression being up- or down-regulated by various abiotic factors (heavy metal ions, drought, salinity, temperature, oxidative stress) and biotic factors (pathogens, insects, fungi) (reviewed by **Bai et al., 2023**), no studies have yet examined the expression patterns of jute LAC under stress conditions. The expression patterns of the *CcaLAC28* and *CcaLAC32* (homologous to *AtLAC17*) genes were validated under two different stress conditions: ABA hormone stress and heavy-metal copper stress.

Why were these (copper and ABA) specific abiotic stresses chosen? Copper is an essential heavy metal and a cofactor for laccase, playing a critical role in its catalytic function. The GO term assignment revealed that the *CcaLAC* genes are associated with the molecular function of copper ion binding (GO:0005507) (**Supplementary Figure 6**). This suggests that some of the identified 32 *CcaLAC* genes might respond to copper, a hypothesis that was further tested under 0.10 mM CuSO_4_·H_2_O stress. Under copper stress, significant differences in *LAC* gene expression were observed, with *CcaLAC28* showing a 23.96-fold increase at 12 h and *CcaLAC32* showing a 45.73-fold increase at 8 h compared to the 0 h samples (**Figure 6A**). Such high expression was also previously observed in rice laccase 10 (*OsLAC10*) under copper stress, with a 1200-fold increase at 12 h (**Liu et al., 2017**). Our goal was to identify *CcaLAC* genes involved in lignification, and we applied copper stress, as reported previously, to identify responsive genes that could be laccases. Further biochemical analysis will be required to confirm their laccase activity. It should be noted that, while the expression changes reported here were obtained empirically from stress-treated plants rather than predicted *in silico*, transcriptional responsiveness alone does not establish a functional or adaptive role for *CcaLAC28* and *CcaLAC32* in stress tolerance; direct measurement of laccase enzymatic activity and lignin content under stress conditions would be needed to establish this link.

ABA hormone stress mimics multiple stress conditions in plants, such as desiccation, freezing, and high salinity. Additionally, literature supports that drought stress can alter laccase expression in plants (**Sharma et al., 2023**). We found that *CcaLAC28* and *CcaLAC32* (homologous to *AtLAC17*) responded to ABA stress (**Figure 6B**). The presence of cis-acting elements such as MYC, MYB, CAT-box, and W-box, which are known for ABA responsiveness in plants, likely explains this expression pattern (**Fujita et al., 2011**). These elements are also involved in drought and salinity stresses, suggesting that these genes might respond to those conditions as well. Further studies on the *CcaLAC* gene expression and function under various biotic and abiotic stresses will provide more information on LAC’s upstream cis-acting elements and their interactions with stress.

The expression of *CcaLAC*s in plant cells (subcellular localisation) was predicted *in silico* (**Supplementary Table 4**). Nucleus-specific localisation was predicted for five *CcaLAC*s (*CcaLAC26, CcaLAC31,* and *CcaLAC32*), whereas nucleus and cytoplasm-specific localisation were predicted for *CcaLAC12* and *CcaLAC34*. *In silico* subcellular localisation studies for *CcaLAC*s provided vital information by predicting their locations in cellular organelles, including chloroplasts, the plasma membrane, endoplasmic reticulum, mitochondria, Golgi apparatus, peroxisomes, vacuoles, the cytoskeleton, and extracellular spaces. The distribution of *CcaLACs* in cells was inferred from *in silico* subcellular localisation data. The presence of *CcaLACs* in all these membrane-bound compartments could be attributed to the presence of transmembrane helices in the majority (18 of 32) of *CcaLAC*s, as found in our analysis (**Supplementary Figure 1**). The structure of the plant laccase is still not well studied. Information about the presence of transmembrane helices could provide valuable insights into its structure.

Functional role analysis of *CcaLAC* genes through Gene Ontology (GO) term assignment identified 32 *CcaLAC* genes in white jute as being associated with key biological processes, including the phenylpropanoid biosynthetic (GO:0009699) and metabolic (GO:0009698) pathways (**Supplementary Figure 6**). Lignin, a crucial structural polymer in plants, is a product of the phenylpropanoid pathway. The GO term analysis further predicted that these *CcaLAC* genes are involved in lignin biosynthesis (GO:0009809) and metabolism (GO:0009808). While the GO term assignment supports these functional roles, experimental validation is required to confirm these predictions.

Manipulation of *LAC* genes has successfully altered lignin content in several crops. Downregulation of *GhLAC1* in cotton, *TaLAC4* in wheat, and overexpression of LACs in pear and silvergrass (*Miscanthus* sp.) significantly impacted lignin deposition (**Hu et al., 2020; Cheng et al., 2019; He et al., 2019**). In jute, targeted downregulation of lignin pathway genes has already been achieved, and phloem-specific silencing using the AtSUC2 promoter offers a strategy to reduce lignin without compromising plant fitness (**Majumder et al., 2020b**). The present findings are correlative, derived from expression, homology, and *in silico* structural/interaction evidence; they do not constitute direct functional proof. Given their expression patterns and regulatory features, *CcaLAC28* and *CcaLAC32* emerge as promising targets for developing low-lignin jute varieties and warrant detailed functional validation, which is currently in progress in our laboratory using stable transformation approaches. Given their expression patterns and regulatory features, *CcaLAC28* and *CcaLAC32* emerge as promising targets for developing low-lignin jute varieties and warrant detailed functional validation.

## 5. Conclusion

The white jute (*C. capsularis*) genome harbours 32 putative laccase genes distributed across its seven chromosomes. The expression of these genes was validated using publicly available transcriptomic datasets, revealing that a majority are preferentially expressed in phloem tissue. Gene Ontology (GO) annotation further supports their involvement in lignin biosynthesis and associated phenylpropanoid pathways. Among the identified laccases, *CcaLAC28* and *CcaLAC32* emerged as strong candidates for roles in stem lignification, based on their close sequence homology to well-characterised Arabidopsis lignin-associated laccases. It should be emphasised that these associations are based on expression, homology, and structural evidence rather than direct functional assays; knockdown or overexpression studies, currently underway, will be required to establish a causal role. Both genes exhibited a progressive increase in expression during plant development but showed significantly reduced expression in the low-lignin mutant *dlpf*, which contains approximately 50% less lignin than the control. Collectively, these findings provide a comprehensive foundation for understanding laccase-mediated lignification in white jute. Future functional studies employing gene editing, transgenic approaches, or reverse genetics will be required to establish the precise roles of *CcaLAC28* and *CcaLAC32* in lignification, ultimately fulfilling the industrial demand for low-lignin jute fibre for diversified biodegradable products.

## Acknowledgments

We extend our sincere gratitude to the Director of the ICAR-Central Research Institute for Jute and Allied Fibres (CRIJAF), Kolkata, India, for providing the jute seeds. We also thank Prof. Swapan K. Datta and Dr Karabi Datta from the Department of Botany, University of Calcutta, Kolkata, India, for their valuable suggestions. We are deeply appreciative of the late Dr Ajay Parida, former Director, and Dr Debasis Dash, Director of ILS, Bhubaneswar, Odisha, India, for their unwavering support and essential facilities critical to the success of this project. We acknowledge Mrs Soma Roy for her valuable comments and suggestions as an internal reviewer.

## CRediT Authorship Contribution Statement

**Subhadarshini Parida:** Formal analysis, Investigation, Methodology, Writing – original draft. **Deepak Kumar Jha:** Formal analysis, Investigation, Methodology, Writing – original draft. **Khushbu Kumari:** Formal analysis, Investigation, Methodology, Writing – original draft. **Seema Pradhan:** Formal analysis, Data curation, Methodology, Writing– review & editing. **Nrisingha Dey:** Methodology, Writing - review & editing. **Shuvobrata Majumder:** Conceptualisation, Investigation, Formal analysis, Funding acquisition, Methodology, Writing - review & editing.

## Data availability

Data is provided within the manuscript or supplementary information files. A preprint version of this article is available at https://doi.org/10.1101/2024.07.17.603856

---

**SUPPLEMENTARY FIGURE 1:** Prediction of transmembrane helices in CcaLACs.

**SUPPLEMENTARY FIGURE 2:** Cis-acting elements in the upstream regions of CcaLACs.

**SUPPLEMENTARY FIGURE 3:** Validation of *in silico* heatmap (transcriptomics) data using qRT-PCR.

**SUPPLEMENTARY FIGURE 4:** Comparative study of laccase (*CcaLAC*) gene expression in white jute cultivar JRC212 and its mutant line *dlpf*. This figure presents a comparative expression analysis of laccase (*CcaLAC*) genes in JRC212 and *dlpf* (which has fibres with 50% lower lignin content) across different developmental stages (30 DAS to 120 DAS).

**SUPPLEMENTARY FIGURE 5:** MicroRNA target prediction in *CcaLAC*s

**SUPPLEMENTARY FIGURE 6:** Gene Ontology (GO) term classification of the *CcaLAC* gene family.

**SUPPLEMENTARY TABLE 1**: List of primers

**SUPPLEMENTARY TABLE 2**: List of orthologous and paralogous pairs of *CcaLAC* genes

**SUPPLEMENTARY TABLE 3**: Bootstrap values of phylogenetic clades

**SUPPLEMENTARY TABLE 4**: Protein features and chromosomal details of identified 32 *CcaLAC*s members in *Corchorus capsularis*

**SUPPLEMENTARY TABLE 5**: Ka/Ks Values

**SUPPLEMENTARY TABLE 6**: Functions of identified cis-regulatory elements in *CcaLAC* genes

**SUPPLEMENTARY TABLE 7**: miRNA Targets in *CcaLAC*s

