## Supplementary figures and images for "Genome-wide Identification of the Laccase Gene Family in White Jute (*Corchorus capsularis*): Potential Targets for Lignin Engineering in Bast Fibre"

### Supplementary Figure 1

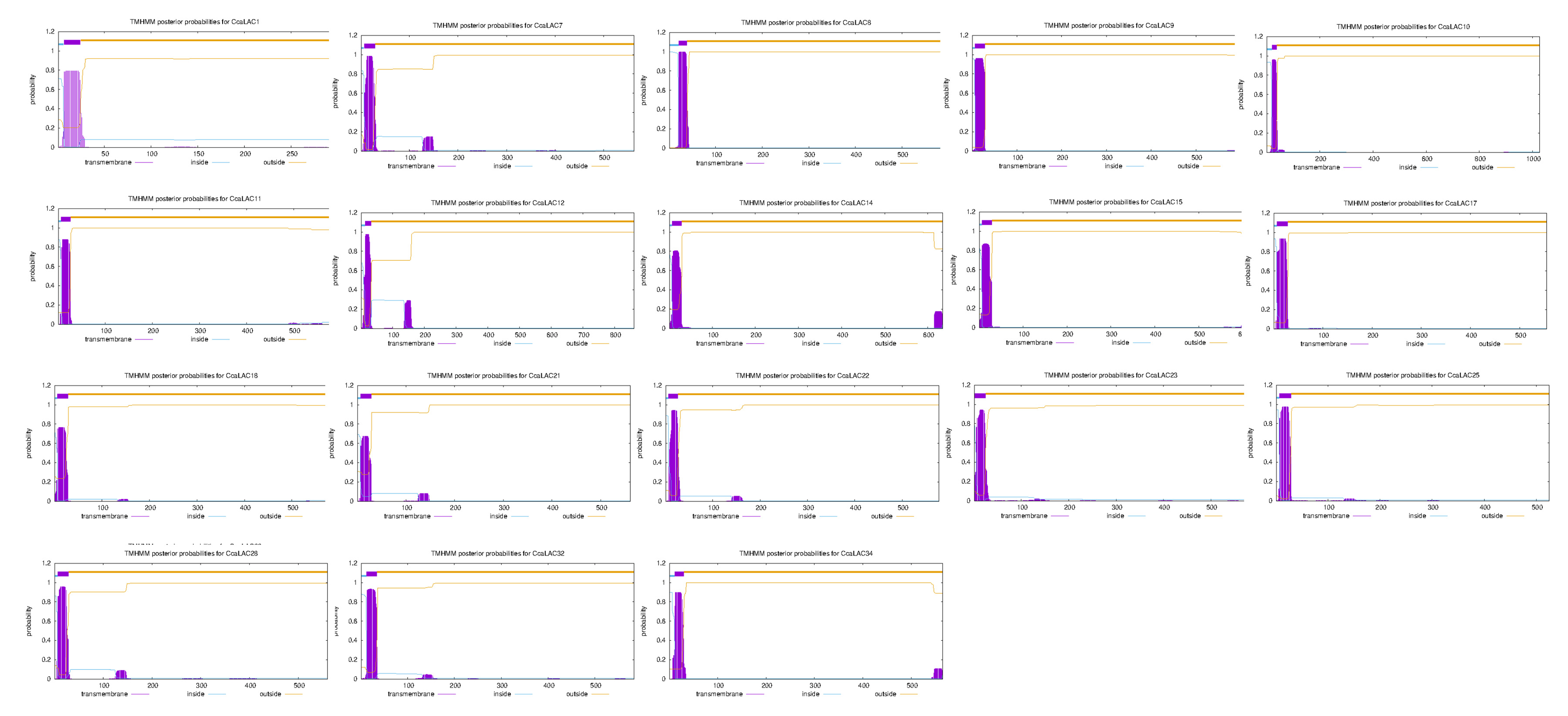

### Supplementary Figure 2

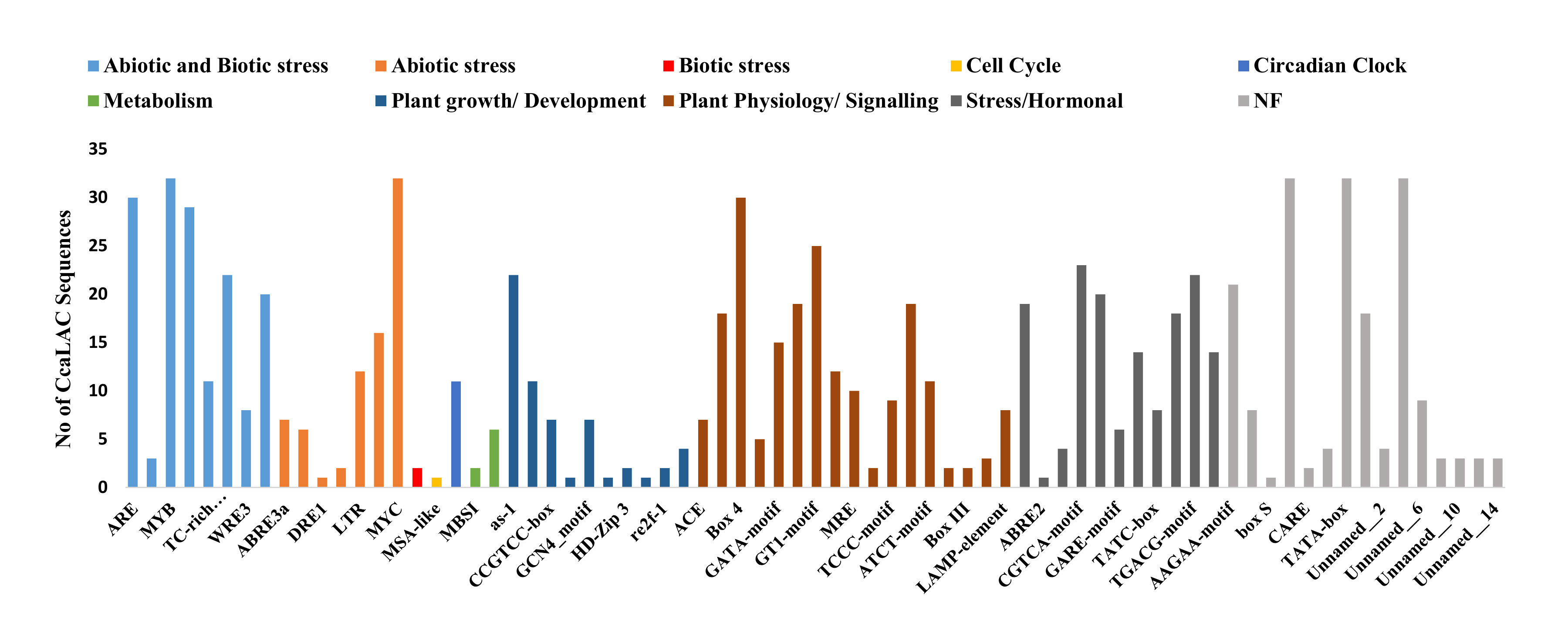

### Supplementary Figure 3

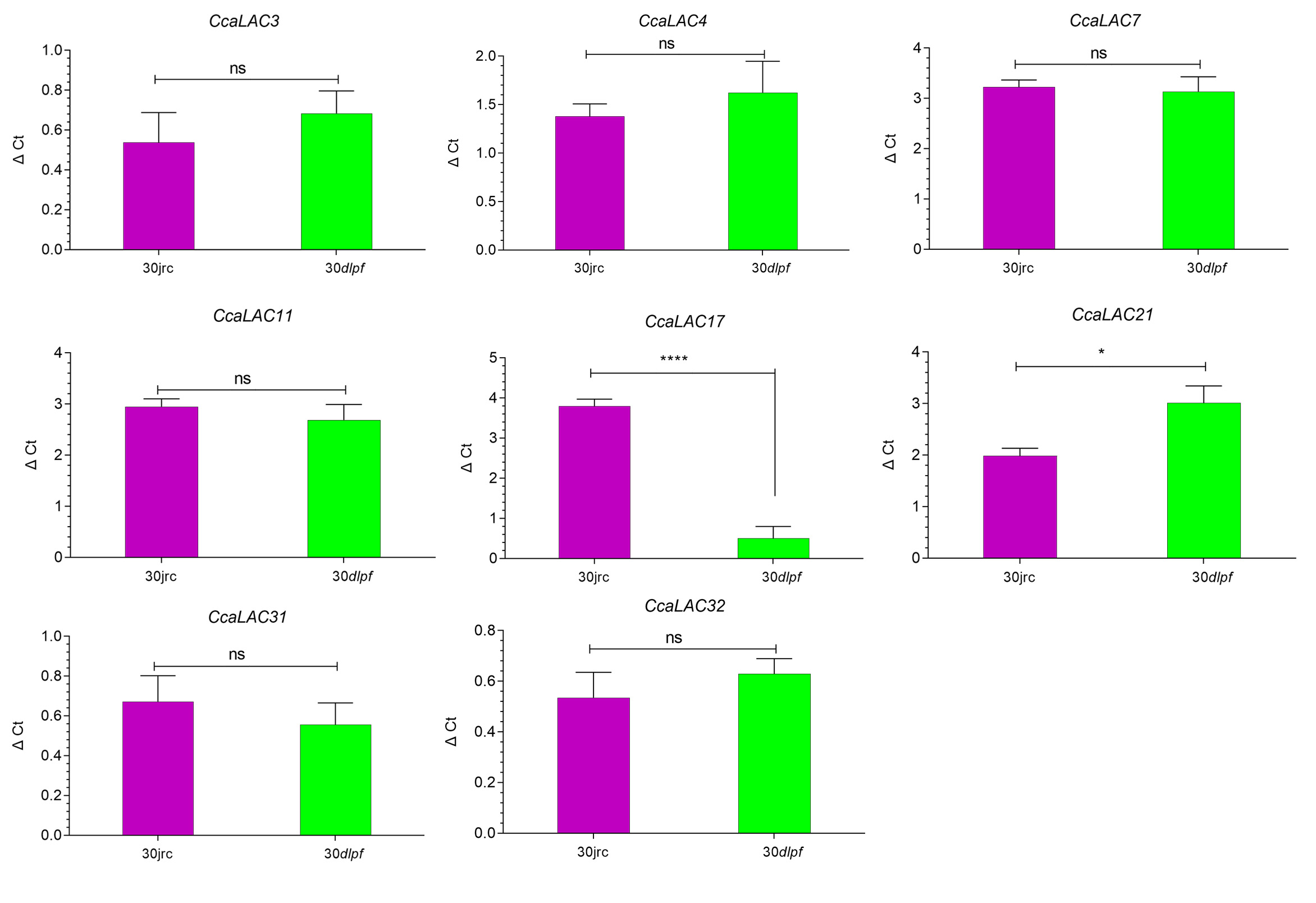

### Supplementary Figure 4

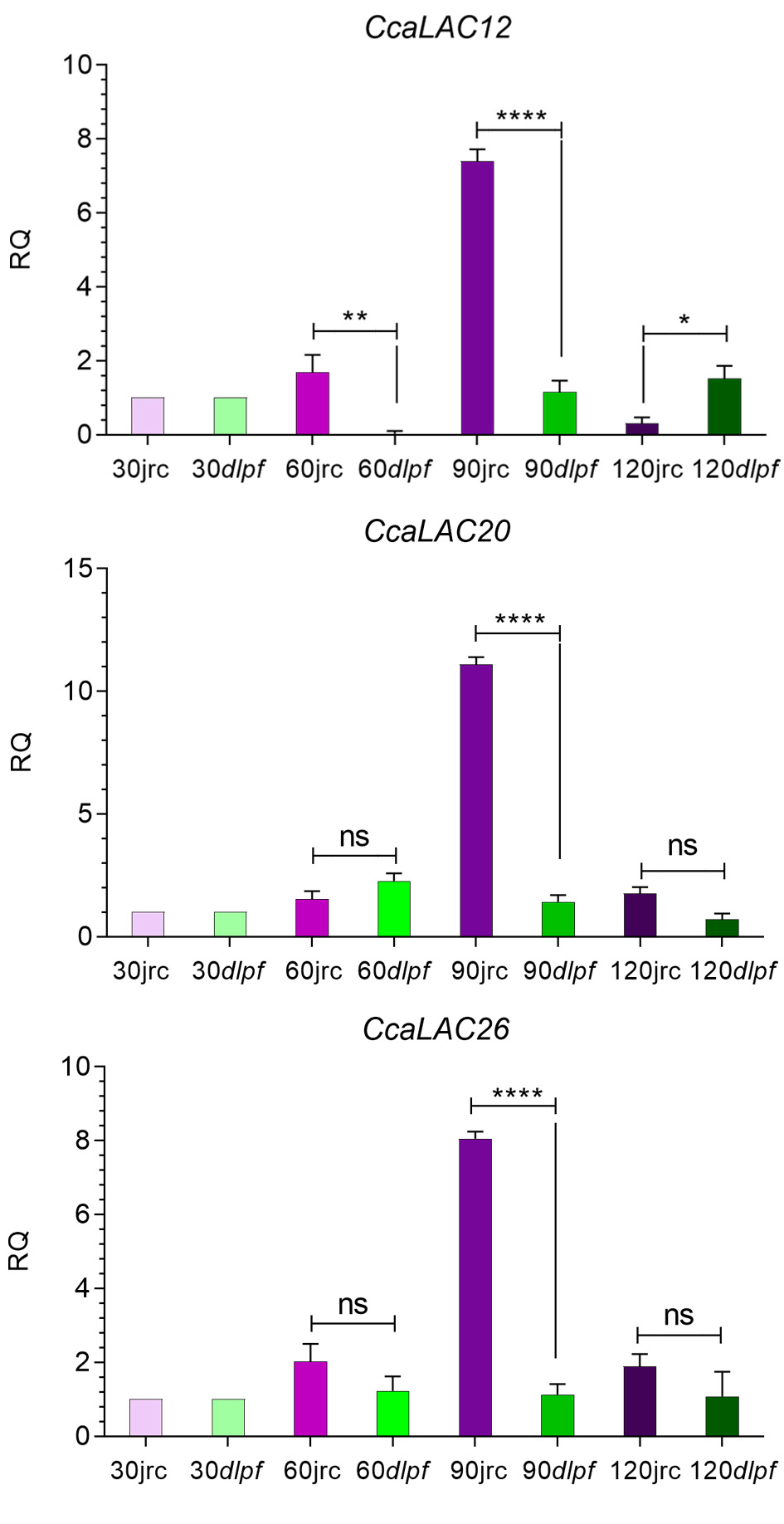

### Supplementary Figure 5

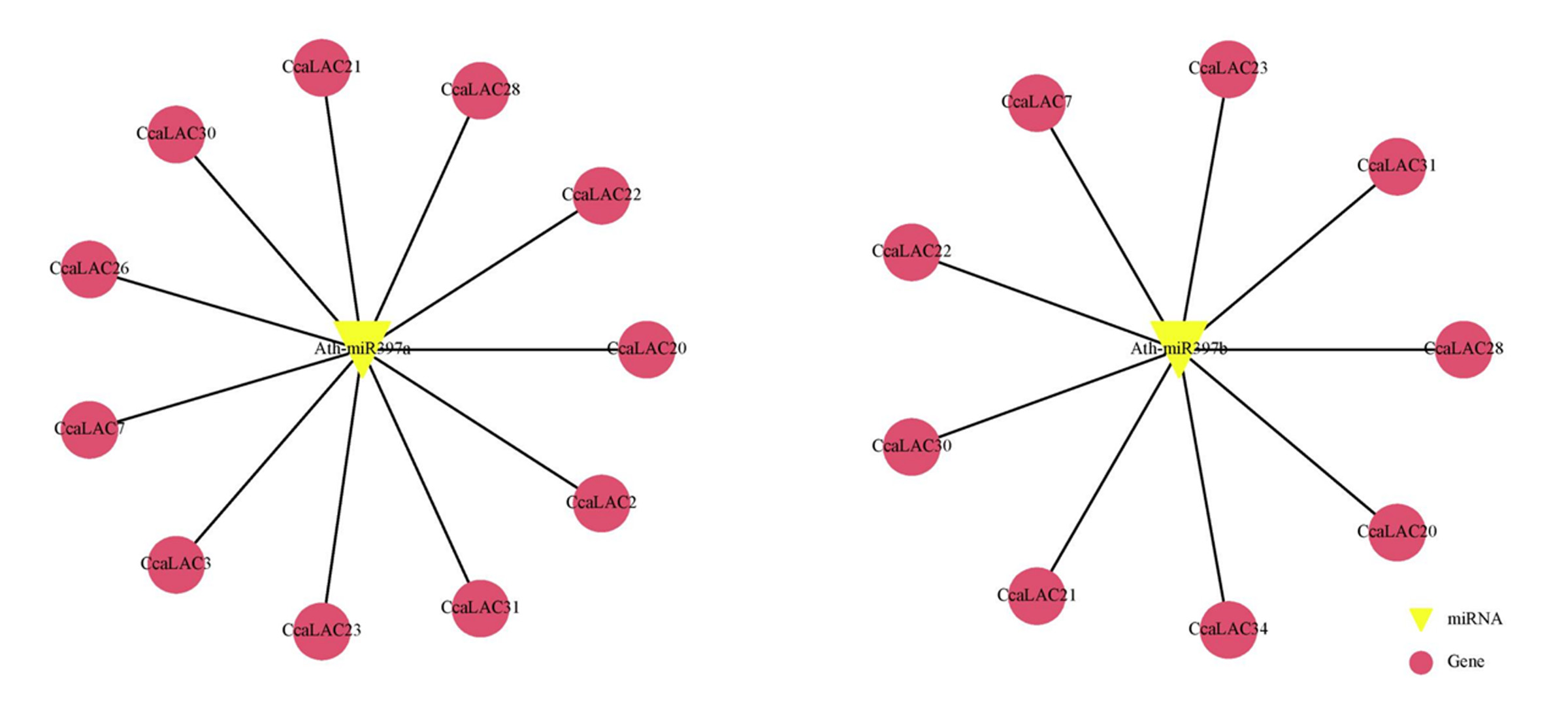

### Supplementary Figure 6

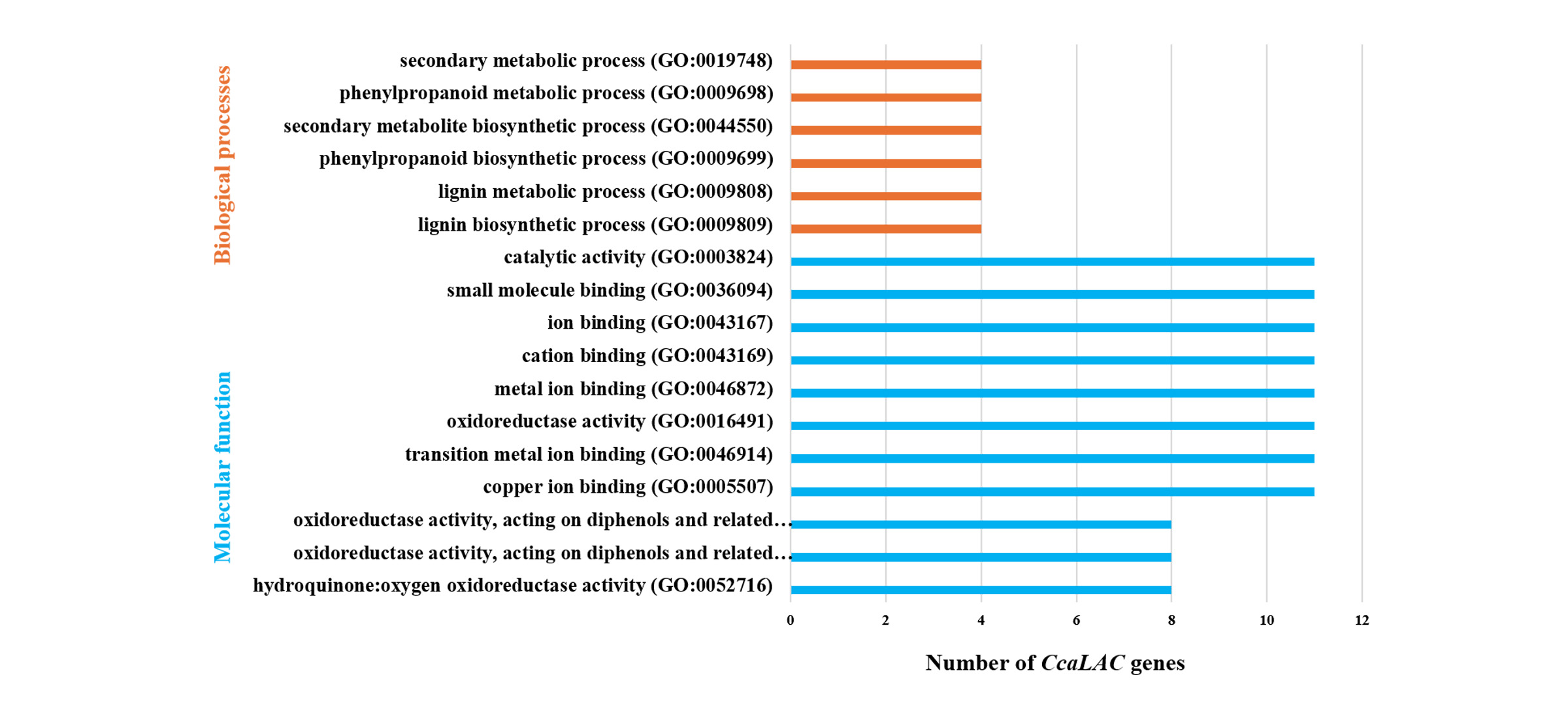
